# LINE-1 activation in the cerebellum drives ataxia

**DOI:** 10.1101/2021.12.03.471151

**Authors:** Takehiro Takahashi, Eriko Kudo, Eric Song, Fernando Carvalho, Yuki Yasumoto, Yong Kong, Annsea Park, Milan Stoiljkovic, Xiao-Bing Gao, Marya Shanabrough, Klara Szigeti-Buck, Zhong-Wu Liu, Yalan Zhang, Parker Sulkowski, Peter M. Glazer, Leonard K. Kaczmarek, Tamas L. Horvath, Akiko Iwasaki

## Abstract

Previous studies have revealed that dysregulation of long interspersed nuclear element 1 (LINE-1), a dominant class of transposable elements in the human genome, correlates with neurodegeneration^1–3^. Yet whether LINE-1 dysregulation is causal to disease pathogenesis has not been proven directly. Here, we demonstrate that expression of evolutionarily younger LINE-1 families is elevated in the cerebella of ataxia telangiectasia (AT) patients, which was correlated with extensive downregulation of epigenetic silencers. To examine whether LINE-1 activation causes neurologic disease, we established an approach to directly target and activate the promoter of a young family of LINE-1 in mice. LINE-1 activation in the cerebellum was sufficient to lead to robust progressive ataxia. Purkinje cells in the diseased mice exhibited marked electrophysiological dysfunctions and degeneration with a significant accumulation of cytoplasmic ribonucleoprotein LINE-1Orf1p aggregates, endoplasmic reticulum (ER) stress, and DNA damage. Treatment with lamivudine, a nucleoside reverse transcriptase inhibitor, blunted the disease progression by reducing DNA damage, attenuating gliosis and interferon gene signature, and recovering the loss of key functional molecules for calcium homeostasis in Purkinje cells. This study provides direct evidence that young LINE-1 activation drives ataxia phenotype, and points to its pleiotropic effects leading to DNA damage, inflammation, and dysfunction and degeneration of neurons.

## INTRODUCTION

The long interspersed nuclear element 1 (LINE-1, L1) is an autonomously replicating genomic parasite that has been amplifying in mammalian genomes for more than 100 million years. LINE-1 constitutes ∼20% of mammalian genomes, and is the only actively replicating retroelement in the human genome^4^. Mounting evidence shows aberrantly high expression and activities of LINE-1 in the context of neurological disorders suggesting the roles of LINE-1 in their etiopathogenesis^1–3^. Studies using knockout mice or cell-based studies using patient-derived cells deficient in the regulators of LINE-1 have shed light on the mechanisms by which aberrant activities of LINE-1 can disturb myriad of physiological processes of the host. These include deficiencies in nucleases that degrade LINE-1 cDNA, TREX1, or RNA, ADAR and RNAaseH2; as well as the reader of methylated DNA, MeCP2, and mediators of DNA repair such as TDP-43, SIRT6, and ATM^1–3^.

A recent transcriptome study of transposable element (TE) expression in the developing human brain^5^ demonstrated that cerebellum exceptionally retains high expression of key epigenetic silencers of TEs such as TRIM28 and DNMT1, both of which have critical roles in regulating young LINE-1^6, 7^, throughout fetal development and even after birth. In contrast to the cerebellum, these TE silencers are significantly downregulated in all other brain regions after birth. Although this evidence suggested distinct expression pattern of TE regulators in the cerebellum, the relevance for this specific brain region to maintain higher expression of these regulators has not been explored. Here we analyzed expression of LINE-1 in the cerebella from patients with three most prevalent types of hereditary ataxia. In parallel, to probe the impact of LINE-1 activation in the cerebellum, we constructed a new mouse model with cerebellar LINE-1 overexpression based on CRISPR approach, and investigated LINE-1 activation-associated phenotype development.

### Loss of key TE silencers and young LINE-1 activation in ataxia telangiectasia cerebellum

First, we examined postmortem samples of cerebellar vermis from patients diagnosed with hereditary cerebellar ataxia of the following types: Ataxia Telangiectasia (AT), Spinocerebellar ataxia type 3 (SCA3) and Friedreich’s ataxia patients, along with the non-affected individuals (information in Extended Data Table 1). Analysis of mRNA expression revealed significant upregulation of LINE-1, coupled with a marked downregulation of key epigenetic silencers *TRIM28* and *DNMT1* in AT compared to unaffected individuals (Fig. 1a). Interferon stimulated genes (ISGs) were significantly upregulated in AT (Fig. 1b) and showed strong correlation with LINE-1 mRNA expression (Extended Data Fig. 1a), suggesting the role of LINE-1 in neuroinflammation^1–3, 8^. Using RNA samples from AT patients and age-/sex-matched controls, we performed RNA sequencing (n = 6 on 6, information in Extended Data Table 1). Principal component analysis (PCA) showed clear segregation of two groups in terms of cellular genes and TE expression (Extended Data Fig. 1b). We found marked upregulation of ISGs and pathways belonging to “inflammatory response” and “defense response to virus” in line with qRT-PCR results (Fig. 1c, gene ontology (GO) analysis of differentially expressed genes (DEGs) in Extended Data Fig. 1c). Significantly upregulated genes included astrocyte marker *GFAP*, possibly indicating the astrogliosis in AT cerebellum (Fig. 1c). The most downregulated genes included *ITPR1*, *CALB1*, *CALB2*, *PVALB* genes (Fig. 1c), encoding inositol 1,4,5-trisphosphate receptor type 1 (IP3R1), Calbindin, Calretinin, and Parvalbumin, respectively. These are all well-known cellular markers of Purkinje cells as well as the critical regulators of calcium ion homeostasis in these cells, and their downregulations have been associated with hereditary ataxias such as SCA1, 15, 16, 29 (Ref. ^9, 10^). AT patients had generally higher expression of LINE-1 across many subfamilies (Fig. 1d). However, analysis at the family level revealed that significant differences were selectively found in younger LINE-1 families, i.e., L1HS (human), L1PA2-17 and L1PB (primate), in contrast to the lack of differences in more ancient families such as L1MA/ME (mammalian) (Fig. 1e). Endogenous retroviruses (ERVs) were also upregulated in AT. However, clustering between patients and controls was more distinct for LINE-1s than ERVs (Fig. 1d and Extended Data Fig. 1d). We also noticed extensive downregulation of critical regulators of TEs in AT cerebella (Extended Data Fig. 1e and 1f), including TRIM28, nucleosome remodeling and deacetylase (NuRD) complex, Human Silencing Hub (HUSH) complex, reader of DNA methylation MeCP2, and piwi-interacting RNA (piRNA)- related MOV10L1 (Fig. 1f, Extended Data Fig. 1e and 1f). Of note, the expression levels of these regulators showed robust negative correlation with the younger families of LINE-1 such as L1HS, L1PA, and L1PB, but not with the older families (Fig. 1f. Correlation plots for *TRIM28* and L1HS/L1ME shown in 1g). Thus, a strong negative correlation was found for key TE epigenetic regulators and selective de-repression of the young LINE-1s in AT cerebellum.

**Fig. 1.**
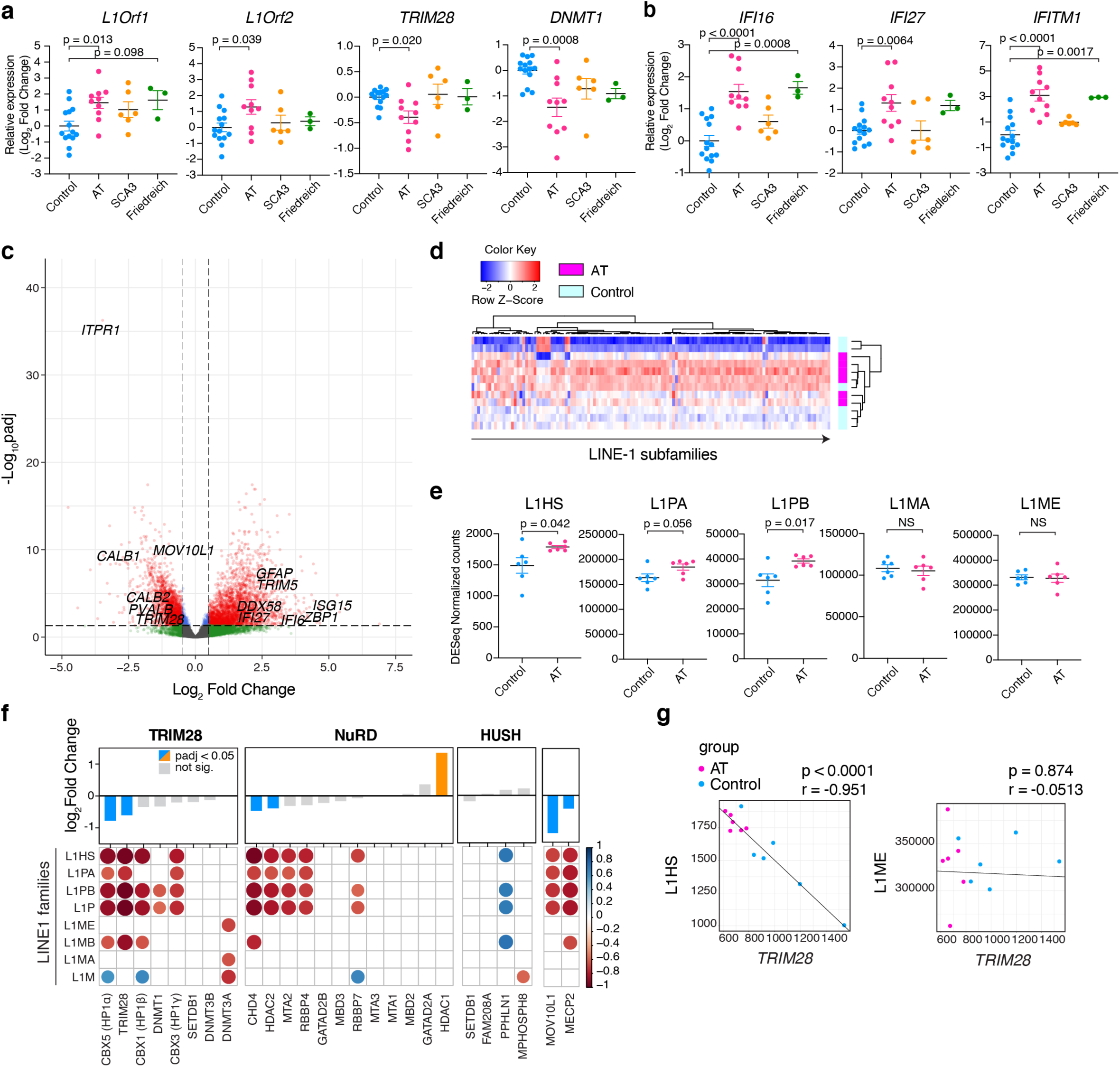
Expression of LINE-1 and its regulators in the cerebellar ataxia patients’ cerebella. **a**, mRNA expressions of ataxia patients and controls. n = 14, 10, 6, and 3 for controls, AT, SCA3, and Friedreich’s ataxia patients, respectively. Data are mean ± SEM. p-values with One-way Anova with Bonferroni’s multiple comparison test for the comparison to control group. Values are shown as log-2 fold change with the average of controls set at 0. **b**, Expression of representative ISGs. **c**, Volcano plot of cellular genes from bulk RNA-seq of AT and control cerebella. n = 6 per group. DESeq2 adjusted p-values and log-2 fold change compared to control samples are plotted. **d**, Heatmap for LINE-1 subfamilies across samples. Each column represents LINE-1 subfamilies. **e**, Comparisons of normalized counts for each LINE-1 family. The counts for subfamilies were added up to calculate the counts per each family (L1HS=L1PA1, L1PA: L1PA2-17, L1PB: L1PB/PB1-4, L1MA: L1MA1-10, L1ME: L1ME1-5/L1MEa-j. p-values shown are with unpaired two-sided t-test. **f**, Summary of correlations between expression (normalized counts) of TE regulators and each LINE-1 family. Representative TE regulators are selected and categorized to TRIM28-, NuRD-, and HUSH-related, according to the list of regulators in a previous literature^50^, and two known regulators of LINE-1, MECP2 and MOV10L1, are additionally included. A bar graph above represents the log2-fold change (AT over control) and adjusted p-values of each regulators, and correlation heatmap below represents the Pearson correlation coefficient between these regulators and each LINE-1 family. **g**, Representative correlation plots from **f**, showing correlations between *TRIM28* and L1HS or L1ME expressions. Pearson correlation coefficients and p-values are shown.

### Establishment of CRISPR activation system of young mouse LINE-1 in vitro

To examine the functional consequences of LINE-1 activation, we set out to establish a system in which young LINE-1 is directly manipulated in mice. We adopted clustered regularly interspaced short palindromic repeat (CRISPR)-mediated approach for future in vivo application. We targeted the Tf family, which is the youngest mouse LINE-1 family^11^. The promoter sequence consists of 7.5 times repeat of 212 bp monomer sequence^12^, and we designed five 20 bp sgRNAs targeting within this monomer sequence, which are not mutually overlapping. We transduced NIH-3T3 cells with dCas9-VP64/MPH CRISPR-activation (CRISPRa) machinery^13^ (hereafter dCasVP-3T3 cells), and transfected Tf-promoter targeting sgRNA- or scramble (Scr) sgRNA-expression plasmid to these cells. Since we are targeting the youngest family of LINE-1, which retains intact protein coding sequence, we assessed the Orfp1 protein expression for the effectiveness of sgRNAs. While Scr_sgRNA and 4 out of 5 promoter targeting sgRNAs led to undetectable or weak Orf1p expression, we found one sgRNA (hereafter L1_sgRNA) induced Orf1p expression which was robustly detectable from 24-hour post transfection (Extended Data Fig. 2a). We decided to use this L1_sgRNA as the LINE-1 targeting sgRNA in vitro as well as in vivo for the mouse generation later in the study. Next, we examined the mRNA expression of L1_sgRNA or Scr_sgRNA expression plasmid-transfected dCasVP-3T3 cells at 48-hour post transfection with qRT-PCR. We found a significant increase in *Orf1* and *Orf2* mRNA expression, which was accompanied by significant increase in the ISG expressions in L1_sgRNA- transfected condition (Extended Data Fig. 2b). Next, we examined the transcriptome of these cells with RNA-seq. We found a significant upregulation of young LINE-1 subfamilies in L1_sgRNA condition, with the most significant increase in the expression of Tf_I subfamily belonging to Tf family (Fig. 2a, left). Other upregulated LINE-1 subfamilies belonged to Tf, Gf, or A families, meaning that young LINE-1 families^11^ were preferentially upregulated, while less differences were detected in older LINE-1 families between scramble and L1_sgRNA conditions (Fig. 2a). We next analyzed cellular gene expression changes. GO analysis of DEGs showed that anti-viral genes and ISGs were the most differentially expressed between these conditions (Fig. 2b, GO terms highlighted in magenta. Heatmap for DEGs in “defense response to virus” term, the top GO term in Fig. 2b, shown as Fig. 2c). Further, we identified multiple terms related to ion homeostasis (Fig. 2b, GO terms highlighted in green, heatmap for DEGs in “ion transport”, the second from the top in Fig. 2b, shown in Extended Data Fig. 2c) and in particular, terms related to cellular calcium homeostasis, which is known to be dysregulated in the context of viral infections^14^.

**Fig. 2.**
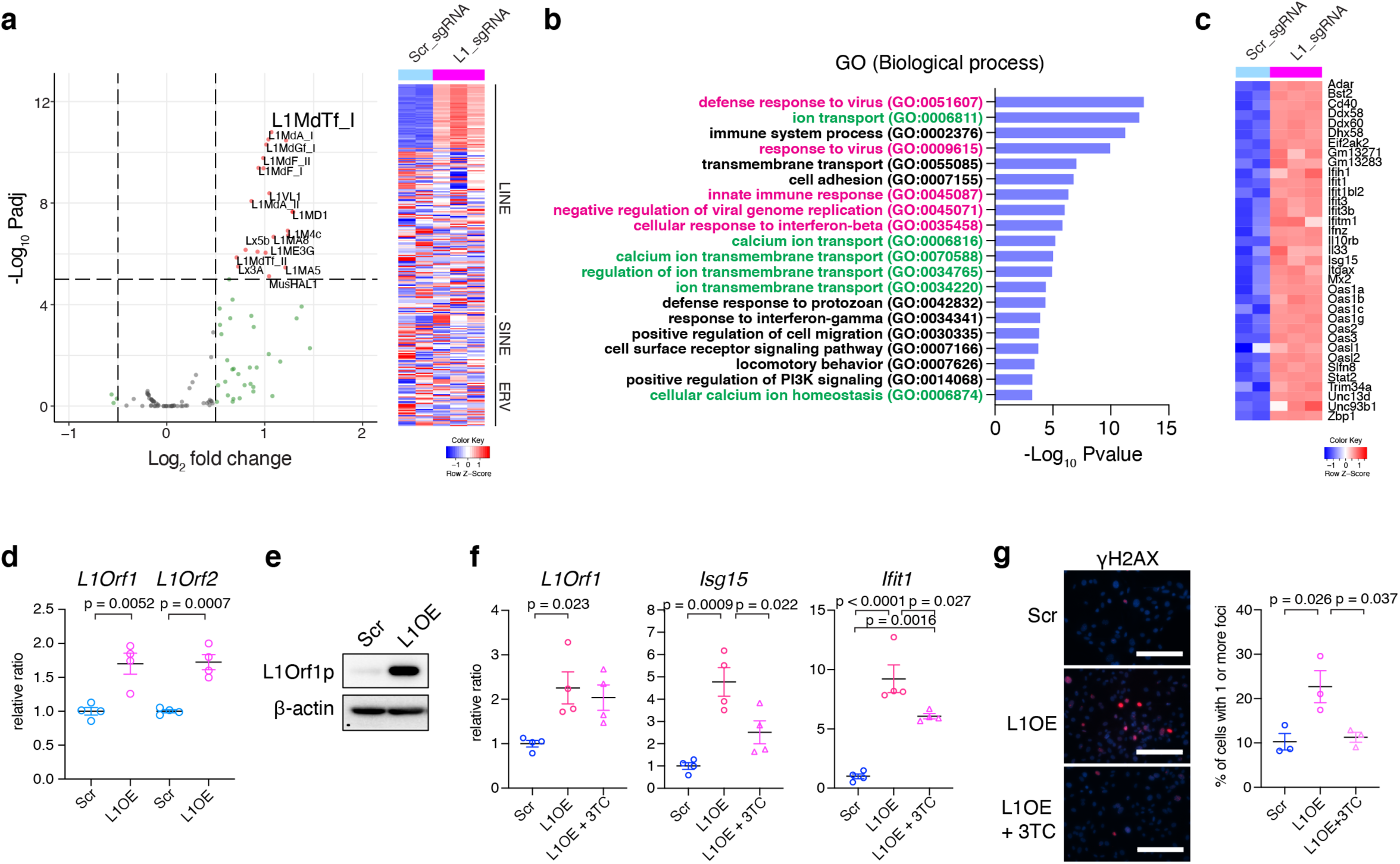
Establishment of LINE-1 CRISPRa system in vitro. **a**, RNA-seq was performed using total RNAs extracted from dCasVP-3T3 cells transfected with Scr_sgRNA or L1_sgRNA (48- hour post-transfection, n = 2 and 3, respectively). Fold changes and adjusted P values for each LINE-1 subfamily plotted as a volcano plot (Left). Heatmap of the TEs across the samples. Each row represents TE subfamily. ERVs are uniquely mappable ERVs in mice^36, 37^. **b**, Gene Ontology analysis (Biological Process) of the DEGs. Top 20 GO terms are shown. Anti-viral/type-I interferon response-related terms and ion homeostasis-related terms are highlighted in magenta and green, respectively. **c**, Heatmap of DEGs in the “defense response to virus” category (the top GO term in **b**). **d,** LINE-1 CRISPRa (LINE-1 overexpression; L1OE) in primary MEFs. MEFs were transduced with dCas9-VP64, MPH, and sgRNA-lentivector, selected with antibiotics for 96 hours, passaged, and 48 hours later, mRNA and protein expression were assayed. mRNA expression levels were normalized to *Gapdh* expression. n = 4. **e**, Whole cell lysates of the same conditions were assayed for LINE-1Orf1p protein expression. **f**, L1OE MEFs were treated with 3TC or DMSO and incubated for 48 hours. Control cells (scramble condition) were treated with DMSO as well. L1Orf1 and ISG mRNA expression were assayed. n = 4 per condition. **g**, The L1OE cells and control cells treated either with 3TC or DMSO were stained for γH2AX. Representative images are shown (left). The foci number were counted. More than 100 cells were analyzed per independent replicate, and % of cells with more than one focus are shown. Result from 3 independent replicates is summarized (right). Scale bar = 200 μm. Data are mean ± SEM. p-values with unpaired two-sided t-test in **d** and one-way Anova with Tukey’s post-hoc test in **f,** and **g**.

We further applied this LINE-1 CRISPRa system to primary cells. Stable transduction of MPH, dCas9-VP64, and L1_sgRNA in primary mouse embryonic fibroblasts (MEFs) led to increased LINE-1 mRNA expression, ISG induction, and LINE-1 protein expression compared to the control, MPH, dCas9-VP64, and Scr_sgRNA transduced MEFs (L1 overexpression/OE MEFs and Scr MEFs, respectively. Fig. 2d and 2e). Previous studies have shown that the nucleoside reverse transcriptase inhibitor (NRTI), lamivudine (3TC), suppresses retrotransposition and ISG induction by LINE-1^2, 8, 15, 16^. We examined if the phenotypes of LINE-1 overexpression in MEFs can be rescued by 3TC treatment. Despite little impact on LINE-1 mRNA expression level as expected, 3TC treatment significantly reduced ISGs (Fig. 2f). LINE-1 retrotransposition activities are known to cause DNA damage^17^, and we found increased γH2AX foci in the nuclei of LINE-1 CRISPRa MEFs, which were reduced by 3TC treatment (Fig. 2g). Thus, we established a system in which young mouse LINE-1 is selectively activated by its direct targeting, and provided evidence of ISG induction, dysregulation in genes involved in ion homeostasis, and DNA damage.

### LINE-1 activation in the cerebellum leads to ataxia in vivo

Next, we sought to generate an in vivo model in which LINE-1 is directly activated by crossing dCas9-CRISPR activator transgenic (Tg) mice with Tg mice for L1 sgRNA. We first obtained dCas9-SPH Tg mice (hereafter dCas mice), which have a transgene encoding dCas9 and CRISPR activator SPH under the pCAG promoter followed by Lox-Stop-Lox (LSL) sequence^18^ (Fig. 3a). We generated two strains of sgRNA Tg mice, namely, Scr_sgRNA mice and L1_sgRNA mice (Fig. 3a, Extended Data Fig. 3a). The progenies of both strains were fertile and had normal litter sizes. In order to test the functionality of transgene sgRNA, we established MEFs from these Tg mice and transduced with dCas9-VP64/MPH CRISPRa machinery. We constructed LINE-1 Tf promoter luciferase reporter (L1_Luc reporter) plasmid and transfected this to the transduced cells. MEFs from L1_sgRNA mice showed significantly enhanced luciferase activities compared to the MEFs from Scr_sgRNA mice, demonstrating the functionality of sgRNA in L1_sgRNA mice (Extended Data Fig. 3b). Then we crossed these mice with dCas mouse strain to generate dCas/L1_sgRNA and dCas/Scr_sgRNA double transgenic mice (LINE-1a mice and dCasScr mice, respectively, Fig. 3a). Our plan was to cross these mice with Cre recombinase transgenic mice to induce Cre-dependent expression of LINE-1. However, surprisingly, we noticed a robust ataxia phenotype in LINE-1a mice (without Cre), characterized by abnormal gait, head tilt, and circling behavior (images of hindlimb clasping in Fig. 3b). Assessment of dCas mice revealed robust basal expression of dCas9 even without Cre in the cerebellum (Extended Data Fig. 4a and 4b). Previous studies have documented the lack of stringency of Lox-Stop-Lox cassette and basal expression of the downstream sequence in the absence of Cre^19, 20^. The basal expression of dCas9 in the dCas mouse cerebellum was likely not due to an aberrant recombination of Lox sequences but due to the read-through transcription beyond the poly-adenylation signals in the LSL sequence, because PCR analysis using cerebellar genomic DNA did not show any evidence for recombination (Extended Data Fig. 4c. Relevant findings described in the original paper of this strain generation^18^). Staining for dCas9 in the cerebellum samples from WT (C57BL/6J) and dCas mice revealed robust expression of dCas9 exclusively in the molecular layer, with strong signal in Purkinje cells colocalizing with its marker Calbindin (Extended Data Fig. 4c). Modest dCas9 expression was also seen in Glial fibrillary acidic protein (GFAP) positive cells i.e., astrocytes (Extended Data Fig. 4d). Although the reason for this specific pattern of read-through transcription is unknown and merits future investigation, these findings indicated that this strain stably expresses dCas9 principally in Purkinje cells and to the lesser extent in astrocytes without crossing with Cre-driver strains.

**Fig. 3.**
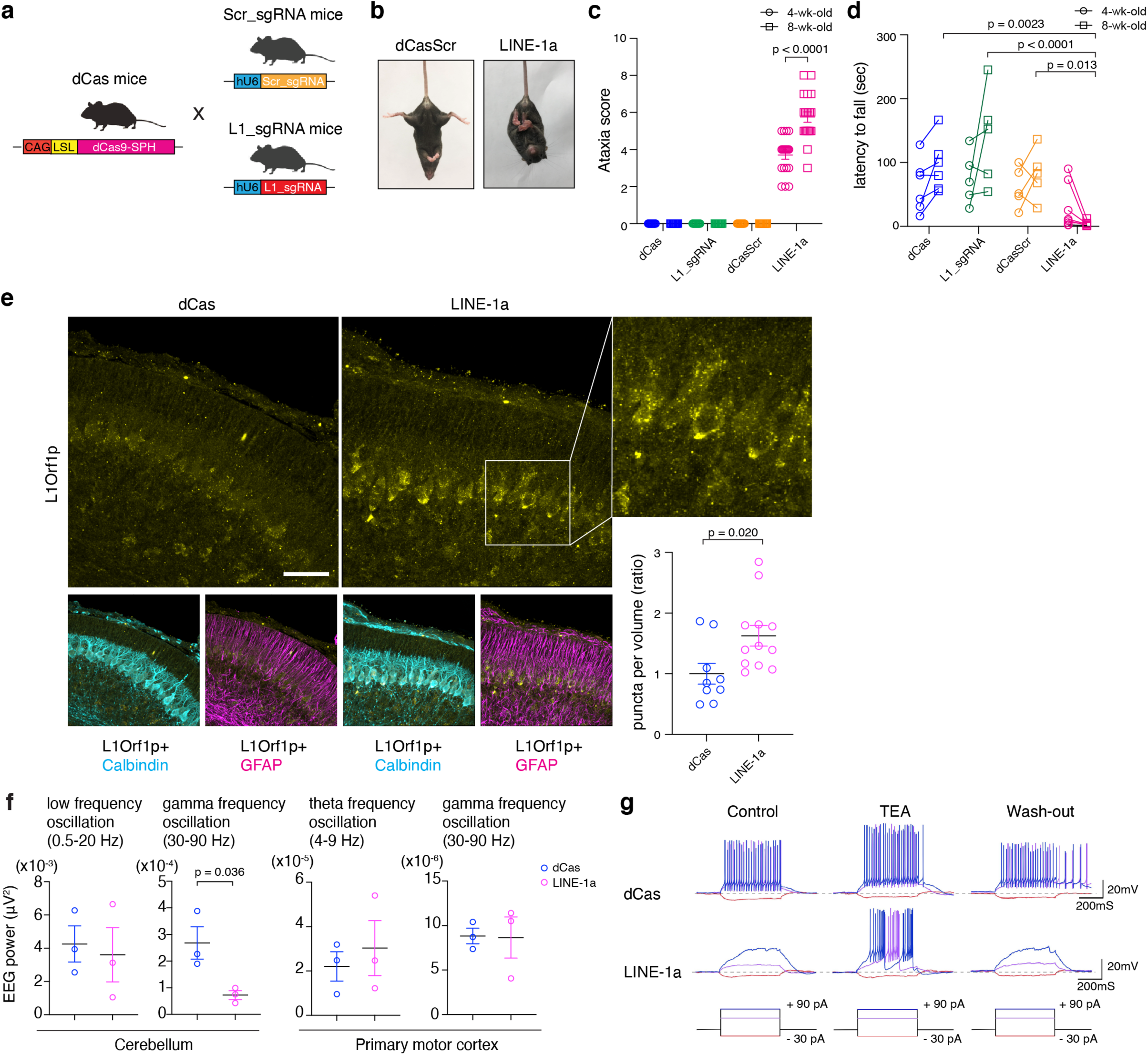
LINE-1 CRISPRa in the cerebellum and development of ataxia. **a**, Transgenic mice harboring Scr_sgRNA or L1_sgRNA were generated (Scr_sgRNA mice and L1_sgRNA mice, respectively), and crossed with dCas9-CRISPR activator (dCas9-SPH) transgenic mice (dCas mice). **b**, LINE-1a mice exhibit hindlimb clasping. Representative images from 8-wk-old dCasScr mouse and LINE-1a mouse. **c**, Composite scores for ataxia were scored at 4- and 8-- wk-old time points. n = 20 (male/female both n=10) for LINE-1a mice. For other genotypes, 10 mice including both male and female were evaluated. **d**, Results of rotarod tests. n = 6, 5, 5, and 8 for dCas, L1_sgRNA, dCasScr, and LINE-1a mice, respectively. **e**, Cerebella from littermate dCas and LINE-1a mice at postnatal day 7 were stained for LINE-1Orf1p, Calbindin, and GFAP. Representative images are shown (Scale bar = 30 μm). Puncta densities inside Purkinje cells per volume were calculated and shown as the ratio (lower right). The individual dot represents data from each 3D field evaluated (n=3 and 4 for dCas and LINE-1a mice, respectively, and images from 3 different lobules in each mouse, n = 9 and 12). p-values with unpaired t-test. **f**, EEG for primary motor cortex and cerebellum in 8-wk-old dCas and LINE-1a mice was recorded and analyzed. n = 3 female mice per group. Similar results were obtained with the same experiment using n = 3 male mice per group. **g**, Representative results of current-clamp electrophysiology study for cerebellar Purkinje cells in the 8-wk-old mice. n = 3 and 4 female mice per group. Seven Purkinje cells per mouse were recorded. The current injection protocols are presented at the bottom. Data are mean ± SEM in **c, d, e, f.**

The ataxia in LINE-1a mice was progressive with age, while none of the littermate dCas mice and L1_sgRNA mice, nor dCasScr mice exhibited this phenotype (Fig. 3c). We also crossed with PcP2 (L7)-Cre mice to further drive Cre expression in Purkinje cells and observed further exacerbated phenotype in these mice compared to double transgenic LINE-1a mice (Extended Data Fig. 5a). There were no statistically significant sex differences in the ataxia severity in LINE-1a mice (Extended Data Fig. 5b). We performed rotarod test, and the latencies to fall were recorded. Average latencies for LINE-1a mice were shorter already at 4-wk-old compared with the age-matched control mice, and at 8-wk of age, LINE-1a mice could hardly keep their body on the rod (Fig. 3d). Since none of the dCas mice, L1_sgRNA mice, dCasScr mice exhibited ataxia phenotype, we used littermate or age- and sex-matched dCas mice as the control group for the LINE-1a group for the subsequent studies.

We examined the cerebella of LINE-1a mice at postnatal day 7 (p7) with immunofluorescence staining and found a significant accumulation of ribonucleoprotein (RNP) LINE1Orf1p in the Purkinje cells compared to the control mice (Fig. 3e). The staining pattern was punctate in the cytoplasm with varying sizes. We assessed spatial density of Orf1p puncta inside the Calbindin staining surface i.e. Purkinje cells, and found significant upregulation of puncta density in LINE-1a mice compared to control mice (Fig. 3e, lower right). We also conducted the staining in 8-wk-old mice. At this age, the staining was seen more diffusely in the entire molecular layer (Extended Data Fig. 5c). We quantified these puncta densities inside the Purkinje cells, and found significantly increased Orf1p puncta in LINE-1a mice (Extended Data Fig. 5d top). We observed the Orf1p staining also within GFAP-positive cells at this age (Extended Data Fig. 5c and 5d bottom). Along with these findings, we noticed markedly disorganized arborization of Purkinje cell dendrites and reduced staining intensities of Calbindin (Extended Data Fig. 5c). Further, enhancement of GFAP staining intensities suggested astrogliosis at 8-wk-old LINE-1a mice (Extended Data Fig. 5c).

Next, we performed electroencephalographic recordings (EEG) in the cerebellum and primary motor cortex (M1). We found a marked decrease in the power of gamma band (30-90 Hz) oscillation in LINE-1a mice in the cerebellum, but no difference in gamma power was detected in M1 (Fig. 3f). Ex vivo current clamp recordings from Purkinje cells from dCas mice showed that supra-threshold current injections induced a train of fast action potentials (APs) in all neurons (7 out of 7 cells from 4 mice) (Fig. 3g, upper left). An application of the potassium channel blocker tetraethylammonium (TEA) at a low concentration (0.5 mM) increased the frequency and the amplitude of APs (Fig. 3g, upper middle), which recovered to the baseline level after washout of TEA (Fig. 3g, upper right). In Purkinje cells from LINE-1a mice, depolarizing currents up to the maximal level applied to the control neurons from dCas animals (90 pA) failed to evoke APs in nearly half of the neurons (3 of 7) from 3 mice cerebella (Fig. 3g, lower left). In these cases, the triggering of APs in response to levels of current that were suprathreshold in control animals, could be reversibly rescued by the application of 0.5 mM TEA (Fig. 3g, lower middle and right). These results collectively demonstrated that Purkinje cells of LINE-1a mice exhibit robust functional alterations resulting in ataxia phenotype development.

### Age-dependent neurodegeneration of Purkinje cells in LINE-1a mice and its attenuation by NRTI treatment

We assessed the number of Purkinje cells in the control and diseased mice. We found that 8-wk-old dCas mice and LINE-1a mice had comparable number of Purkinje cells, while these cells significantly decreased in LINE-1a at the age of 24 weeks compared to controls (Fig. 4a). Electron microscopy (EM) examination of LINE-1a mouse cerebella at 8-week of age revealed the shrinkage of Purkinje cells. They were noticeably darker in color suggesting the high electron density, which had been also reported in Purkinje cells of *Atm* knockout mouse^21^ (Fig. 4b). Numerous dilated ERs in LINE-1a Purkinje cells’ cytoplasm were present, which are characteristic of ER stress (Fig. 4b). Indeed, in these cells, we found significantly upregulated expression of immunoglobulin heavy chain-binding protein (BiP)/glucose-regulated protein 78 (GRP78), the master regulator and marker for unfolded protein responses (UPR) in ER stress (Fig. 4c).

**Fig. 4.**
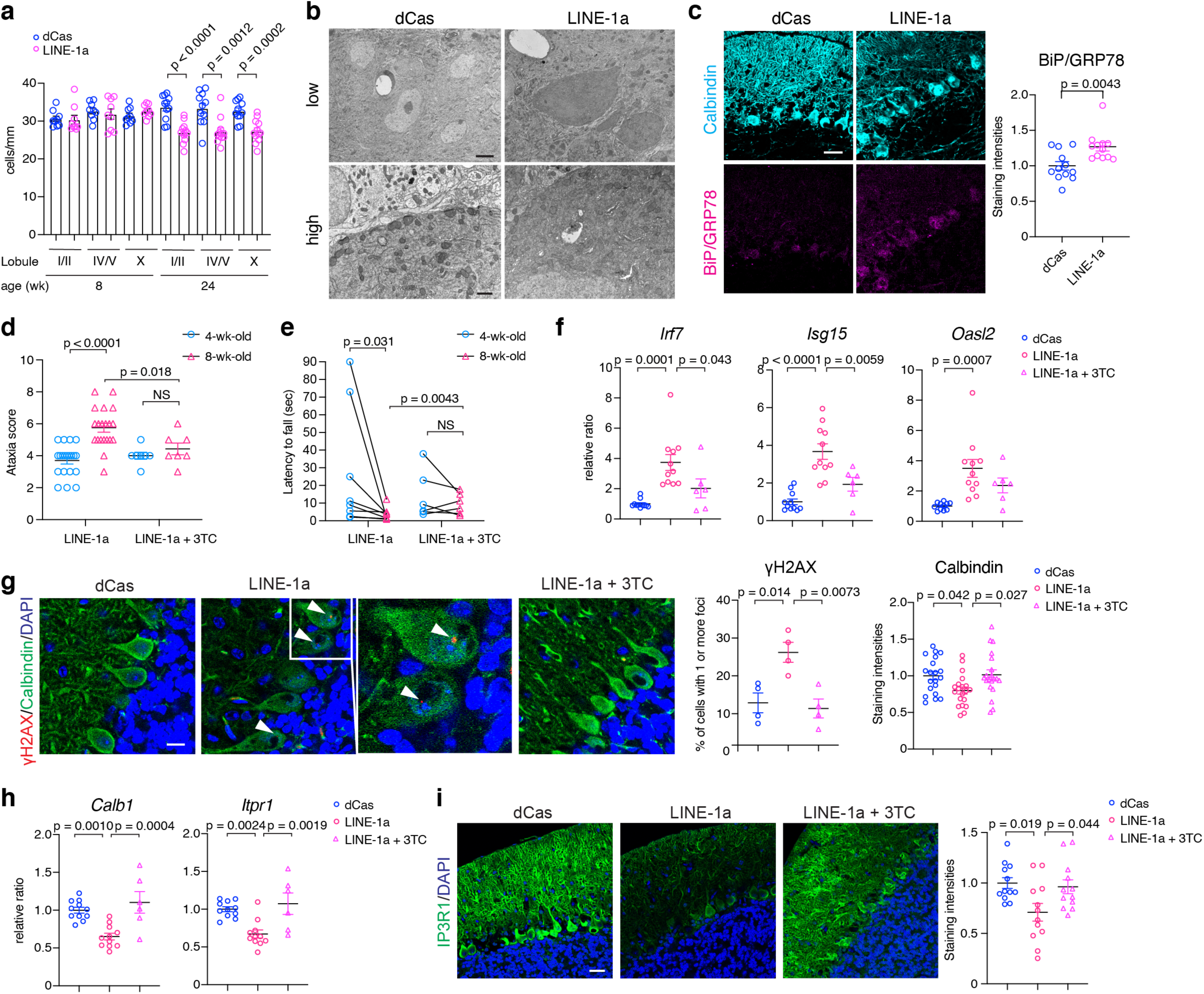
Characterization of LINE-1a mice cerebellum and blockade of phenotype progression with NRTI. **a,** Purkinje cell numbers were counted and normalized to the length. n = 3 or 4 female mice at 8-wk- and 24-wk-old were assessed per group. Three separate sections per mouse were assessed. Each dot represents the number in each section. **b**, Representative electron microscopy images from 8-wk-old dCas and LINE-1a mouse cerebella. n = 3 mice per group. Scale bar = 2 μm for the upper panels, and 500 nm for the lower panels. **c**, Representative immunofluorescence staining of Calbindin and BiP/GRP78 in the cerebellum. Scale bar = 30 μm. Staining intensities of BiP in Purkinje cells were assessed (n = 4 mice per group and 3 independent lobules per mice). **d,** Ataxia scores of at 4- and 8-wk-old naïve LINE-1a mice (n=20, same as in Fig. 3c) and 3TC-treated LINE-1a mice (n=7). Scores of 4- and 8-wk- old mice within the same group are compared with Wilcoxon rank test, and scores across groups are compared with unpaired t-test. **e,** Rotarod test results at 4- and 8-wk-old. n = 8 for naïve LINE-1a (same as in Fig. 3d) and n = 7 for 3TC-treatment group, as in **d**. **f**, ISG expressions were compared between 8-wk-old dCas mice, naïve and 3TC-treated LINE-1a mice (n = 11, 11, and 6, respectively). **g,** Cerebellum in 8-wk-old mice in the same condition as **f** were stained for γH2AX, Calbindin, and DAPI (Scale bar = 10 μm). γH2AX foci in Purkinje cell nuclei were assessed and quantified (middle panel). n = 4 mice per group and more than 50 Purkinje cell nuclei were examined per mouse. Arrow heads: γH2AX foci. Calbindin staining intensities in the same staining were evaluated (right panel). n = 20 fields in dCas, LINE-1a, and 3TC-treated LINE-1a mice, respectively (n=4 mice per group and 5 independent fields per mouse). **h**, mRNA expression of *Calb1* and *Itpr1* in bulk cerebellum (same samples as in **f**). **i,** Representative IP3R1 staining images. Quantification of staining intensities on the right (n = 4 mice per group and 3 independent lobules per mice). Data are mean ± SEM. p-values with unpaired two-sided t-test in **a** and **c** and one-way Anova with Tukey’s post-hoc test in **f, g, h,** and **i**.

Next, bearing in mind the phenotype attenuation by NRTI in vitro (Fig. 2f and 2g), we tested the impact of NRTI treatment in vivo. Following treatment of 4-week-old LINE-1a mice with 3TC (200 mg/day daily by oral gavage) over a period of 4 weeks, we found a significant attenuation of ataxia progression in the treated mice compared to the untreated mice (Fig. 4d and 4e). In line with this, we observed significantly higher levels of ISG mRNA expression in LINE-1a cerebella, which were reduced upon 3TC treatment (Fig. 4f). In parallel, we found a significant decrease in the GFAP staining intensities after 3TC treatment, suggesting attenuated astrogliosis in the treated LINE-1a mice (Extended Data Fig. 6a). Next, we assessed γH2AX foci in Purkinje cell nuclei and found significantly increased number of γH2AX-positive nuclei in LINE-1a mice Purkinje cells, which was reduced by 3TC treatment (Fig. 4g). At the same time, we noticed that the staining intensity of Calbindin that was noticeably reduced in LINE-1a mice in other staining panels (Fig. 4c and Extended Data Fig. 5c), was recovered in LINE-1a mice upon 3TC treatment (Fig. 4g, quantification in right panel). Calbindin is a key calcium buffering protein in Purkinje cells^22^, and its gene *CALB1* was among the most significantly downregulated genes in the AT transcriptome profile (Fig. 1c). The recovery of Calbindin levels was additionally confirmed at the mRNA transcript level as well (Fig. 4h). We also assessed *Itpr1* gene encoding IP3R1, which was the most significantly downregulated gene in the entire AT cerebellum transcriptome (Fig. 1c). IP3R1 is a calcium release channel on ER critically regulating the cellular calcium levels^9, 10, 23^, and heterozygosity of *ITPR1* is causative for hereditary progressive ataxia SCA15/29^9^. In parallel with the changes in Calbindin, we found significant decrease in IP3R1 in LINE-1a mice, which was also recovered by 3TC treatment (Fig. 4h and 4i). Notably, analyzing RNA-seq data from human cerebella, we found that *CALB1* and *ITPR1* expressions strongly correlated with L1HS and *TRIM28* levels (Extended Data Fig. 6b), implicating common molecular pathways of ataxia development in human AT and LINE-1a mice cerebella. Collectively, these findings unmasked that LINE-1 activation in the cerebellum severely damages Purkinje neurons triggering proteostatic and genotoxic stress with concomitant neuroinflammation, whereas NRTI treatment mitigates dysregulation of key functional molecules of calcium ion homeostasis in vivo with attenuated neuroinflammation and DNA damages.

## Discussion

AT is a hereditary cerebellar ataxia caused by mutations in *ATM* gene, a serine/threonine kinase that phosphorylates several key proteins which initiates activation of the DNA damage response, leading to cell cycle arrest, DNA repair or apoptosis. Previous studies have shown that *ATM* regulates the retrotransposition activities of LINE-1^24, 25^. However, whether LINE-1 activation contributes to the pathogenesis of cerebellar neurodegeneration or it is merely a consequence of disease processes remained unclear^24^. Our study showed that the cerebella of AT patients express elevated levels of LINE-1 and inflammatory genes. By generating a mouse model that has elevated expression of LINE-1 in the cerebellum, we demonstrate that LINE-1 expression drives ER stress, DNA damage and inflammation leading to dysfunctional Purkinje cell physiology and degeneration. LINE-1 overexpression was sufficient to drive ataxia phenotype in mice, which was arrested by NRTI treatment. These findings collectively implicate LINE-1 activation as an important pathogenic factor for cerebellar ataxia.

We found severe degeneration of Purkinje cells with cytoplasmic aggregates of ribonucleoprotein Orf1p (Fig. 3e) and alteration in electron density possibly related to ER stress (Fig. 4b and 4c), which is known to disturb cellular calcium homeostasis^26^. Despite the heterogeneity of neurological disorders, a common critical pathological signature is the aggregation of proteins in the cytoplasm, which is associated with ER stress/unfolded protein response (UPR)^27^. Pathological aggregation of RNA-binding proteins/ribonucleoproteins has been demonstrated across various neurodegenerative diseases, and the potential role of LINE-1Orf1p aggregates are suggested in the neurodegeneration of ALS^28, 29^. Although still poorly understood, emerging studies are elucidating the crosstalk between DNA damage/genotoxic stress and proteostatic ER stress responses^27^. Our data suggest that persistent ER stress and DNA damage driven by LINE-1 activation led to electrophysiological disturbances and degeneration of Purkinje cells. Additionally, concomitant glial cell activation and inflammation induced by LINE-1 activation could be playing a role as suggested in *TREX1*-mutant astrocytes which accumulate LINE-1 cDNA in the cytosol^8^.

Our study provides a new model to probe a causal link between LINE-1 activation and cerebellar ataxia. However, we acknowledge several limitations. First, we have generated LINE-1-targeting sgRNA transgenic mice in only one genetic background (C57BL/6). Second, dCas9 basal expression was primarily seen in Purkinje cells but was also present in astrocytes, and LINE-1 overexpression was observed in both cell types (Fig. 3e, Extended Data Fig. 5c), which makes it difficult to assign cell-type specific contribution to the phenotype in LINE-1a mice. Third, we are not able to rule out the off-target CRISPRa effects of non-LINE1 genes. The effects of LINE-1 promoter activation could include the effects on the *cis-*regulatory functions of LINE-1 over gene expressions^30^. Indeed, LINE-1 has been reported to act as the alternative promoters for many genes involved in neuronal functions in neural progenitor cells^31^. LINE-1 activation in the developmental phase may have significant impact in the cerebellar development as well, which cannot be reversed by the treatments. However, the effectiveness of NRTI treatment in blocking the disease progression at the phenotypic as well as at the molecular levels demonstrates that LINE-1 activities in these mice are exacerbating the neurological disease. A previous study has shown that NRTI treatment rescued the tau-induced neurodegeneration in *Drosophila melanogaster* in vivo^32^.

Our collective evidence from LINE-1a mice and AT patient samples implicates the crucial pathological role of LINE-1 activation in cerebellum in the development of ataxia. Our experimental system provides a novel platform to analyze multifaceted consequences of the dysregulation and activation of this parasitic genomic element, and points to the pleiotropic pathological effects caused by TE activation in the neuronal electrophysiological dysfunctions, neuroinflammation, and neurodegeneration.

## Methods

### Mice

C57BL/6J mice (stock #000664) and B6;D2-Gm33925Tn(pb-CAG-cas9*,-EGFP)1Yangh/J mice (dCas mice, stock #031645), B6.129-Tg(Pcp2-cre)2Mpin/J (PcP2Cre mice, stock #004146) were obtained from Jackson. All the mice were bred in-house at Yale University. Mice were housed in SPF condition and care was provided in accordance with Yale Institutional Animal Care and Use Committee guidelines (protocol #10365).

### Plasmids and lentivirus packaging

LentiMPHv2, lenti dCas-VP64_Blast, lentiSAMv2, pX330-U6-Chimeric_BB-CBh-hSpCas9, and sgRNA(MS2) cloning backbone were gifts from Dr. Feng Zhang (Addgene plasmid # 89308 ; http://n2t.net/addgene:89308 ; RRID:Addgene_89308, Addgene plasmid # 61425 ; http://n2t.net/addgene:61425 ;RRID:Addgene_61425, Addgene plasmid # 75112 ; http://n2t.net/addgene:75112 ; RRID:Addgene_75112, Addgene plasmid # 99372 ; http://n2t.net/addgene:99372 ; RRID:Addgene_99372, Addgene plasmid # 42230 ; http://n2t.net/addgene:42230 ; RRID:Addgene_42230, and Addgene plasmid # 61424 ; http://n2t.net/addgene:61424 ; RRID:Addgene_61424, respectively) ^13, 33, 34^. pTN201 was a kind gift from Dr. John Goodier ^12^. The plasmids were amplified in *E. coli* DH5a or Stbl3 (Thermo Fisher) and purified with Plasmid Plus Kit (QIAGEN). The lentiviruses were packaged in Lenti-X 293T cell line (Takara Bio) with Lenti-X packaging single shots (Takara Bio), and concentrated using Lenti-X concentrator (Takara Bio) following the manufacturer’s instructions.

### sgRNA cloning

LentiSAMv2 and sgRNA(MS2) cloning backbone were digested with BsmBI or BbsI (both NEW ENGLAND BIOLABS) in the digestion buffer (NEB 3.1 or 2.1, respectively, NEW ENGLAND BIOLABS) 2 hours at 55 °C or 37 °C, respectively. After digestion, the fragments were electrophoresed in 0.8% agarose gel and purified with Qiaquick Gel Extraction Kit (QIAGEN). Oligos including 20 bp sgRNA sequences plus appropriate overhangs were resuspended in ddH_2_O at 100 μM and annealed in T4 ligase buffer with T4 polynucleotide kinase (both NEW ENGLAND BIOLABS) starting from 95 °C for 5 minutes and then temperature was ramped down to 25 °C at 5 °C/min. The digested vector and annealed oligos were ligated with T4 ligase and T4 ligase buffer (both NEW ENGLAND BIOLABS) at room temperature for 2 hours, transformed in Stbl3 or DH5a competent cells (Thermo Fisher), and plated onto LB plates with appropriate antibiotics. The colonies were picked up for miniprep (QIAprep Spin Miniprep Kit, QIAGEN) and sequencing, and confirmed that the correct sequences were cloned into the sgRNA cloning sites. The sequences of the oligos used for sgRNAs can be found in Extended Data Table 2.

### Generation of L1_sgRNA and Scr_sgRNA Tg mice

L1_sgRNA and Scr_sgRNA Tg mice were designed and generated in collaboration with Yale Genome Editing Center at Yale University. First, pX330 vector (addgene plasmid #42230) was digested with BbsI (NEW ENGLAND BIOLABS), and each sgRNA sequence was cloned into the sgRNA cloning site as described above. The plasmids were amplified in *E. coli* DH5α and purified with a EndoFree Plasmid Maxi Kit (QIAGEN). The plasmids were then digested with AflIII and XbaI (both NEW ENGLAND BIOLABS) to cut out the hU6 promoter and sgRNA sequence. After linearization, the plasmids were microinjected into C57BL/6J x C57BL/6J fertilized oocytes. The incorporation of the transgene into the genome was determined by PCR analysis using genomic DNA extracted from tails. One of the mouse from each strain was chosen as the founder mouse and mated with C57BL/6J mice, and the germ line transmission of the transgene was determined by PCR.

### Mouse LINE-1 Tf promoter reporter plasmid construction

The 5’UTR promoter region of mouse L1Spa belonging to LINE-1 Tf family was amplified with the primers (Fw; AATGGGCAGAGCTCGTTTAG, Rv: CTGGTAATCTCTGGAGTTAG) and Takara LA Taq polymerase with GC buffer (Takara Bio) using pTN201 plasmid (a kind gift from Dr. John Goodier ^12^) as the template. The PCR product was cloned into the pCR Blunt II-TOPO vector using the Zero Blunt TOPO PCR cloning kit (Thermo Fisher). The LINE-1 promoter-luciferase reporter plasmid was constructed by subcloning the promoter fragment into the KpnI-XhoI restriction site of the pGL4.11[luc2P] vector (Promega, Cat No. #E6661). The plasmid was sequenced and verified that the correct sequence was cloned.

### Plasmid transfections, lentiviral transductions, and 3TC treatment

For lentivirus transduction, 2 x 10^5^ NIH-3T3 cells or MEFs were seeded in 6-well cell culture plate. The next day, packaged and concentrated lentivirus supes were added in the cell culture and spinfected for 90 min at 25 °C, 1200 g in the presence of 5 μg/ml polybrene (Santa Cruz). Forty-eight hours after spinfection, antibiotics selection was started. The cells were selected with Hygromycin B (Invivogen) 200 μg/ml, Blasticidin (Thermo Fisher) 5 μg/ml, or Puromycin (Sigma) 2 μg/ml for 4 days. For the analysis of MEFs, MEFs were selected for 4 days with Hygromycin and Blasticidin, passaged, seeded at 4 x 10^4^ cells/well or 2 x 10^5^ cells/well in 24 or 6 well culture dish, respectively, and used for the analyses. For the experiments with 3TC treatments, 3TC obtained from NIH AIDS Research Program was used. Upon passaging after the antibody selection for lentivirus transduction, the treatment with 10 μg/ml 3TC or DMSO was started, and cells and their culture supernatant were collected for mRNA expression/protein expression and ELISA assay 48 hours later, respectively. For plasmid transfection, 4 x 10^4^ cells/well or 2 x 10^5^ cells/well were seeded in 24 or 6 well culture dish, respectively, the day before transfection. The plasmids were transfected to the cells using lipofectamine 3000 transfection reagent (Thermo Fisher). The cells were harvested and analyzed at the time point(s) indicated in each figure legend.

### Reverse transcription-quantitative polymerase chain reaction (RT-PCR)

RNA was isolated from total NIH-3T3 cells, MEFs, mouse tissues, or postmortem human cerebellar vermis samples using either the RNeasy Kit or RNeasy Fibrous Tissue Kit following the manufacturer’s protocols (QIAGEN). For tissue RNA extraction, tissues were harvested into RLT buffer (QIAGEN) and disrupted by the bead homogenization in Lysing Matrix D tubes using a FastPrep-24 5G homogenizer (MP Biomedicals). cDNA was synthesized using Quantitect Reverse Transcription Kit (QIAGEN) from 200 ng isolated mRNA. Quantitative PCR was performed using iTaq Universal SYBR Green Supermix (Biorad). Primers sequences are listed in Extended Data Table 2. For dCas9 mRNA detection, a commercially available specific primer set was used (System Biosciences). For the detection of some other targets, Taqman primer sets were used with TaqMan Fast Advanced Master Mix (ThermoFisher) following the manufacturer’s instructions (Extended Data Table 2).

### RNA sequencing, data processing, and analysis

Two hundred ng of total RNA was used for paired-end library generation with the NEBNext Ultra II RNA Library Prep Kit for Illumina (E7770) for mouse samples. In the library generation of human cerebellum RNA samples, NEBNext rRNA Depletion Kit (NEB E6310) was used. RNA qualities of the human RNA samples (RNA Integrity Number; RIN score) were assayed with high sensitivity RNA Screen Tape (Agilent 5057-5579) and shown in Table S4. Libraries were run on a NextSeq500 to generate 2×75 bp reads for mouse samples and 2x42bp reads for human samples.

Sequencing raw data were aligned using STAR (STAR/2.5.3a-foss-2016b, mm10 assembly for mouse and hg38 for human samples, respectively)^35^ with parameters: --runThreadN 20 --outSAMtype BAM SortedByCoordinate --limitBAMsortRAM 35129075129 -- outFilterMultimapNmax 1 --outFilterMismatchNmax 999 --outFilterMismatchNoverLmax 0.02 -- alignIntronMin 20 --alignIntronMax 1000000 --alignMatesGapMax 1000000 for mapping host genes. For endogenous retroelements, ERVMAP^36, 37^ and RepEnrich2^38^ were used using modules, Python/2.7.13-foss-2016b, Bowtie2/2.2.9-foss-2016b, BEDTools/2.27.1-foss-2016b and SAMtools/1.9-foss-2016b using repeatmasker mm10 assembly. Counts were counted using BEDTools (BEDTools/2.27.1-foss-2016b)^39^, coverageBed function, and normalized using DESeq2 ^40^. The heatmaps and volcano plots were prepared using R version 4.0.5. Gene ontology (GO) analysis were performed with DEGs (genes with |Log_2_FoldChange| > 1, Padj < 1 x 10^-2^), using Database for Annotation Visualization and Integrated Discovery (DAVID) ver 6.8 (NIH/NIAID) ^41^.

### Western Blotting

For cultured cells, cells were lysed with 1x cell lysis buffer (Cell signaling). For mouse tissues, tissues were harvested in lysis buffer (20 mM Tris (pH 7.5), 1% NP-40, 150 mM NaCl with protease inhibitor cocktail (Roche Diagnostics)) and disrupted by the bead homogenization in Lysing Matrix D tubes using a FastPrep-24 5G homogenizer (MP Biomedicals). Lysates were centrifuged at 4°C, 15,000 g for 10 minutes and supernatants were collected. Proteins were fractionated by 10% SDS-PAGE and transferred onto PVDF membranes. The membranes were blocked with TBST with 5% Non-Fat Dry Milk (AmericanBio) for 30 minutes. After blocking, membrane was probed with the following primary antibodies for overnight at 4°C: Rabbit anti-mouse LINE-1Orf1p (Abcam, ab216324, 1:1000), Mouse anti-GAPDH antibody (HRP conjugated, GeneTex, GT239, 1:5000), Rabbit anti-Cas9 antibody (Diagenode, C15310258, 1:1000), Mouse anti-β actin antibody (HRP conjugated, Cell signaling, #5125, 1:2000). After washing, secondary IgG-HRP antibodies (anti-Rabbit IgG, HRP and anti-Mouse IgG, HRP, Cell signaling, #7074 and #7076, respectively. Both 1:1000) were added and incubated for 1 hour at room temperature. The blot was visualized using SuperSignal West Pico PLUS Chemiluminescent Substrate (Thermo Fisher). The band intensities were quantified with ImageJ software (NIH, ImageJ version 1.51m9).

### Generation of mouse embryonic fibroblasts

Pregnant C57BL/6J^-^female mice were sacrificed, and E13.5 or E14.5 embryos were harvested and processed by first removing hearts, livers, and heads. The remaining tissue was placed in a petri dish with 2.5 ml of 0.05% Trypsin-0.5 mM EDTA-PBS and minced with a scissors. The minced tissue was transferred to a 15 mL conical tubes and placed at 4 °C overnight. Then the tissue was incubated in a 37 °C shaking water bath for 30 minutes, and dissociated by adding 3 ml of DMEM with 10% FBS and vigorous pipetting. The isolated cells were filtered through a 70 μm filter, resuspended in 15 ml of DMEM with 10% FBS, and plated in T-175 culture flasks. The cells were grown to confluency prior to freezing down. MEFs were grown in tissue culture dishs/flasks in 10% FBS in DMEM with 1x penicillin-streptomycin.

### DNA damage assessment in vitro

Immunofluorescence staining and analysis for γH2AX foci analysis was performed as previously described ^42, 43^. MEFs were transduced with lentiMPHv2 and lentiSAMv2 with either scr_sgRNA or L1_sgRNA, selected and passaged as described above and treated with 10 μM of 3TC or DMSO for 7 days. Then the cells were trypsinized and seeded in chamber slides. Subsequently, cells were fixed and permeabilized in 1% Formaldehyde/0.2% Triton-X100 for 15 minutes and blocked with 5% bovine serum albumin in PBS for 1 hour at room temperature. Proteins were stained overnight at 4°C with primary antibody (Mouse anti-γH2AX antibody; Millipore Sigma #05-636, 1:300) diluted in blocking solution. Upon washing with 0.5% Triton-X100 in PBS, samples were incubated with secondary antibody (Thermo Fisher, anti-mouse AF 555, 1:1000) for 60 minutes at room temperature. DNA was stained with for 5 minutes at room temperature with DAPI. Data were collected with an EVOS FL microscope (Advanced Microscopy Group). Foci were analyzed with Cell Profiler software ^44^. More than 100 cells analyzed per condition for each experiment.

### Assessment of composite ataxia scores

The mice were assessed on their ataxia score using the composite phenotype scoring system by Guyenet et al.^45^. Mice at the age of 4- and 8-wk-old were evaluated.

### Rotarod test

The mice were placed on a horizontally oriented rod (ACCUROTOE ROTAROD: AccuScan Instruments, Inc) that has an accelerating rotation speed (0-40 rpm over 0-300 seconds). Mice were put on the rod when the rotation speed was 4 rpm, and time to fall off the rod was recorded. Each mouse underwent three trials per day with an interval of 15 minutes between trials. Records for the last performances in the three consecutive trials were used for the analysis.

### *In vitro* electrophysiology

Sagittal cerebellar slices, 300 µm thick, were cut from mice as described previously (Liu et al, 2010). Briefly, dCas and LINE-1a mice were deeply anesthetized with isoflurane and decapitated. Then brains were rapidly removed, trimmed to a tissue block containing only the cerebellum and sectioned with a vibratome in an oxygenated (with 5% CO_2_ and 95% O_2_) cutting solution at 4 °C containing sucrose 220 mM, KCl 2.5 mM, CaCl_2_ 1 mM, MgCl_2_ 6 mM, NaH_2_PO_4_ 1.25 mM, NaHCO_3_ 26 mM, and glucose 10 mM, at pH 7.3 titrated with NaOH. After preparation, slices were maintained in a storage chamber with oxygenated artificial cerebrospinal fluid (ACSF) containing NaCl 124 mM, KCl 3 mM, CaCl_2_ 2 mM, MgCl_2_ 2 mM, NaH_2_PO_4_ 1.23 mM, NaHCO_3_ 26 mM, glucose 10 mM, at pH 7.4 titrated with NaOH. Cerebellar slices were transferred to a recording chamber constantly perfused with ACSF at 33 °C with a rate of 2 ml/min after at least 1-hour recovery.

Whole-cell current clamp was performed to observe membrane and action potential with a Multiclamp 700A amplifier (Axon instrument, CA) as described previously ^46, 47^. The patch pipettes with a tip resistance of 4-6 MΩ were made of borosilicate glass (World Precision Instruments) with a Sutter pipette puller (P-97) and filled with a pipette solution containing K-gluconate 135 mM, MgCl_2_ 2 mM, HEPES 10 mM, EGTA 1.1 mM, Mg-ATP 2 mM, Na_2_- phosphocreatine 10 mM, and Na_2_-GTP 0.3 mM, at pH 7.3 titrated with KOH. After a giga-ohm (GΩ) seal and whole-cell access were achieved, the series resistance (between 10 and 20 MΩ) was partially compensated by the amplifier. To test the action potential firing properties in Purkinje cells, a series of current injections (initial = -30 pA, increment = 20 pA, duration 500 ms, 7 steps) were applied to recorded cells. However, the exact intensity of current injection was adjusted according to input resistance in each recorded neuron.

All data were sampled at 10 kHz and filtered at 3 kHz with an Apple Macintosh computer using AxoGraph X (AxoGraph, Inc.). Electrophysiological data were analyzed with AxoGraph X and plotted with Igor Pro software (WaveMetrics, Lake Oswego, OR).

### *In vivo* electrophysiology

Local field potential (LFP) recordings were performed *in vivo* from the primary motor cortex (M1) and cerebellum in dCas and LINE-1a mice (n=3 for each genotype and each sex) using a 16- channel silicon probe (A1x16; NeuroNexus Technologies, Inc). Eight-week-old mice were anesthetized with urethane (1.5 g/kg) given by intraperitoneal injection, and placed in a stereotaxic frame (Kopf) on a temperature-regulated heating pad (Physitemp Instruments Inc.) set to maintain body temperature at 37-38 °C. After unilateral craniotomies above the targeted regions, recording probe was positioned and lowered first in the M1 region (1.1 mm anterior from bregma, 1.5 mm lateral, and 1.0 mm ventral from the brain surface) and then in the cerebellum (6.2 mm posterior from bregma, at midline and 1.7 mm ventral from the brain surface) using coordinates from the stereotaxic atlas ^48^. Before the beginning of recordings at each site, animals were allowed to stabilize for 15 minutes, whereupon spontaneous LFPs from M1 and cerebellum were measured over 20 minutes. Recording signal was amplified using A-M System amplifier (model 3600, Carlsborg) with filters set between 1 Hz and 500 Hz, digitized at a rate of 1 kHz, and collected on a computer via a CED Micro1401-3 interface and Spike2 software (Cambridge Electronic Design).

For quantitative offline analyses, LFPs were subjected to Fast Fourier transform at a spectral resolution of 0.24 Hz using Spike2. The range of the bands/oscillations are defined as follows: Theta band: 4-9 Hz, Gamma band: 30-90 Hz, and low oscillation (encompassing delta, theta, alfa and low beta) from 0.5-20 Hz. The magnitude of power of spontaneous oscillatory activity was computed in 5-minute long epochs of stable LFP signals for each animal by averaging power in a given band across the 5 superficial channels (15-11) for both M1 and cerebellum. For determining gamma oscillation power, the signal was first band-pass filtered between 30 Hz and 90 Hz. All values were initially determined to be suitable for parametric analysis according to normality and homoscedasticity.

### Purkinje cell count and measurement

Purkinje cell numbers were counted using 3 different sections of midsagittal cerebellar sections for each mouse. Cells in each lobule were counted, and normalized to the length measured with ImageJ.

### Immunohistochemistry and immunofluorescence staining of the mouse cerebellum

For immunohistochemistry, mice were euthanized and perfused transcardially with 4% PFA in PBS, pH 7.4. Brains were kept in 4% PFA in PBS overnight at 4 °C, dehydrated in 70 % ethanol, and embedded in paraffin. Sections were made and slides were stained for Hematoxylin and Eosin (H&E) at the Yale Pathology Tissue Services Core.

For immunofluorescence studies, mice were euthanized and perfused transcardially with 4% PFA in PBS, pH 7.4. Brains were kept in fixation solution overnight at 4 °C, transferred to 30% sucrose solution for 24 hours, and embedded in O.C.T. (Tissue-Tek) for frozen sectioning in a cryostat. Cerebellar parasagittal sections were cut at a thickness of 40 μm through the vermis. Tissues were permeabilized, washed and stained while free-floating in 24-well cell culture plate.

First, tissues were permeabilized with 0.3% Triton X-100 in PBS, then incubated with 10% normal donkey serum (Jackson ImmunoResearch) in 0.3% Triton X-100 in PBS (blocking solution) for 1 hour, and subsequently immunoreacted with the primary antibodies (Mouse anti-Calbindin antibody, Abcam, ab82812, 1:200; Chicken anti-GFAP antibody, Thermo Fisher, PA1-10004, 1:200; Rabbit anti-mouse LINE-1 Orf1p antibody, Abcam, ab216324, 1:100, Rabbit anti-IP3 Receptor 1 antibody, Thermo Fisher, PA1-901, 1:200, Rabbit anti-γH2AX antibody, Cell Signaling, #9718, 1:200, Rabbit anti-BiP antibody, Cell signaling, #3177, 1:100, Rabbit anti-Cas9 antibody, Cell signaling, #19526, 1:100, anti-GFP antibody, abcam, ab290, 1:100) at 4 °C for 16 hours. Tissues were washed with PBS, immunoreacted with secondary antibodies (Goat anti-Mouse IgG Alexa Fluor 488, Goat anti-Chicken IgY Alexa Fluor 555, Donkey anti-Rabbit IgG Alexa Fluor 647, all from Thermo Fisher and used at 1:1000 dilution), washed with PBS, stained with DAPI (Sigma-Millipore), washed with PBS, and tissues were placed on the slide glasses. Coverslips were mounted with ProLong Gold Antifade mountant (Invitrogen). Z-stack images (1μm step) from three different cerebellar lobules (lobule I/II, IX, and X) per mouse were acquired using the same settings on Leica SP8 on HC PL APO 40x/1.30 oil-immersion. Images were analyzed for staining surface, staining intensities, γH2AX foci, and LINE-1Orf1p puncta with the image analyzer IMARIS (Imaris 9.2, Bitplane AG). The puncta were quantified using spot detection function of the software (expected puncta radius set at 1 μm). The number of puncta inside Calbindin staining surface (Purkinje cells) or GFAP staining (astrocytes) surface were divided by the volume inside each surface to calculate the puncta densities per volume in each cell type.

### DNA damage assessment in cerebellar Purkinje cell nuclei in vivo

For the assessment of γH2AX foci in Purkinje cell nuclei, γH2AX, Calbindin, and DAPI were co-stained in cerebellum as described above, and images were obtained with Leica SP8 on HC PL APO 60x/1.30 oil-immersion. Each visual field contained 7-15 Purkinje cell nuclei, and presence of γH2AX foci inside each Purkinje cell nucleus was evaluated. Foci inside nuclei that were visible in two consecutive Z-stack images (Z-step 1 μm) were counted as γH2AX foci. Images from 4 to 6 fields from at least 3 different lobules were collected in order to assess > 50 Purkinje cell nuclei per mouse, and percentage of foci positive Purkinje nuclei among the total nuclei assessed were calculated for each mouse.

### Electron microscopy

Mice were deeply anesthetized and were perfused with Somogyi-Takagi fixative ^49^. The cerebellum was cut with Vibratome in 50 μm thick sections. After osmication, the sections were dehydrated in ascending ethanol and propylene oxide, and flat-embedded and polymerized in Durcupan ACM (EMS14040). 70 nm ultrathin sections were cut on Leica ultramicrotome, and the images were collected on transmission electron microscope (FEI). The analysis was carried out in a blinded manner.

### 3TC treatment of the mice

Lamivudine (Selleckchem, S1706) was dissolved in PBS at the concentration of 2 mg/ml, and 100 μl solution was given to the mice with oral gavage daily (200 mg/day) for 4 weeks, starting from 4-week of age and up to 8-wk of age. On the next day of the final treatment, the mice were evaluated for ataxia scoring and rotarod tests, euthanized, and the cerebella were collected for experiments.

### Data availability

Data deposition: The RNA-seq data in this paper will be deposited in the Gene Expression Omnibus (GEO) database.

### Statistical analysis

Statistical analyses were performed using GraphPad Prism 8. The reported values are means ± SEM, as noted in the figure legend for each panel. Statistical significance was determined using One-way Anova with Bonferroni’s multiple comparison test, one-way ANOVA with Tukey’s multiple comparison test, unpaired two-sided t-test, or Wilcoxon matched-pairs signed rank test, as designated in the figure legends. A p-value of < 0.05 was considered to be statistically significant.

## ACKNOWLEDGEMENTS

We thank Melissa Linehan and Huiping Dong for technical and logistical assistance. We thank Dr. William Philbrick and Dr. Timothy Nottoli at Yale Genome Editing Center for the assistance in generating the transgenic mice. We thank NIH NeuroBioBank from which we obtained all the human frozen brain samples. We thank Dr. John Goodier at Johns Hopkins School of Medicine for providing us the pTN201 plasmid used in the generation of the mouse LINE1 promoter luciferase reporter. We thank Dr. Alex Bortvin at Carnegie Institution of Science for the protein extract of Maelstrom knockout mouse testis and LINE1Orf1p antibody used for the validation of our LINE-1Orf1p detection. We want to thank all current and former members of the Iwasaki lab for insightful discussions. This work was supported by the Howard Hughes Medical Institute (A.I.), National Institutes of Health (NIH) grants R01 NS111242 (A.I. and L.K.K.), R01 AG067329, R01 052005 and P01 AG051459 (T.L.H.), the Uehara Memorial Foundation (T.T.), and the Japan Society for the Promotion of Science (T.T.).

## AUTHOR CONTRIBUTIONS

Conceptualization, T.T., A.I., T.L.H., L.K.K.; Investigation, T.T., E.K., F.C., E.S., A.P., Y.K., Y.Y., M.S., X.G., M.S., K.S-B., Z.L., Y.Z., P.S.; Writing, T.T., A.I.; Supervision, A.I., T.L.H., L.K.K., P.M.G.; Funding Acquisition, A.I., T.L.H., L.K.K.

---

**Extended Data Fig. 1.**
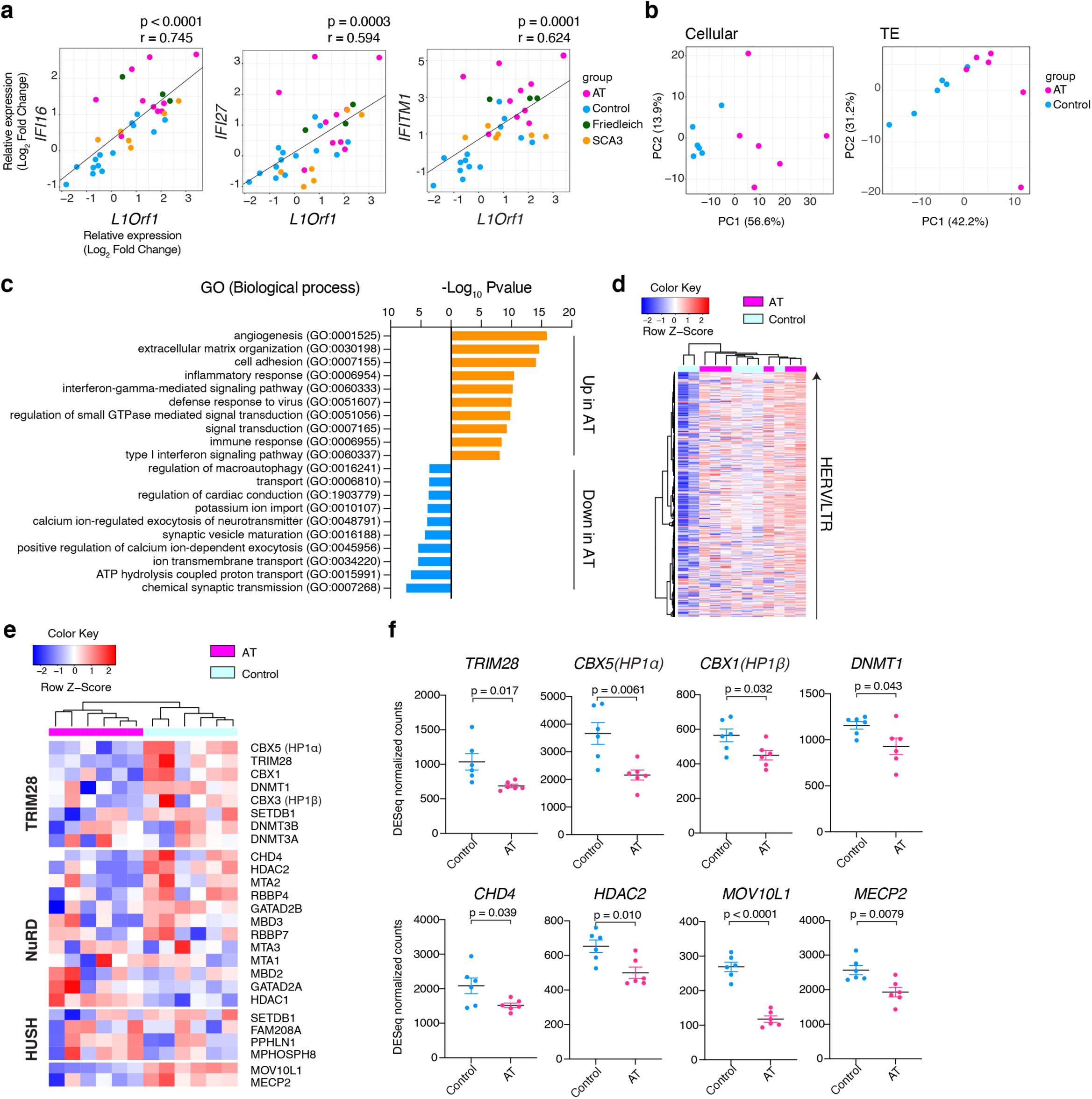
Transcriptomic feature and downregulation of TE regulators in cerebellar ataxia patients. **a**, Correlation of L1Orf1 and ISG mRNA expressions. Pearson correlation (r) and p-values for the correlations are indicated above each panel. Values are shown as the log-2 fold changes from the mean values of control group as in Fig. 1a. **b**, PCA plot for cellular gene expression (left) and TE expression (right) with RNA-seq data from control individuals and AT patients (n = 6). **c,** GO term analysis (Biological Process) of DEGs between AT and controls. DEGs (padj < 0.01, |Log2-fold change|>1) are used for the analysis. Top-10 GO terms for upregulated and downregulated genes are shown. **d,** Heatmap of ERVs in AT and controls. Each row represents one ERV or LTR family. **e**, Heatmap for representative TE regulators . **f**, DESeq normalized counts are shown for 8 TE regulators. Data are mean ± SEM. p-values with unpaired two-sided t-test are shown.

**Extended Data Fig. 2.**
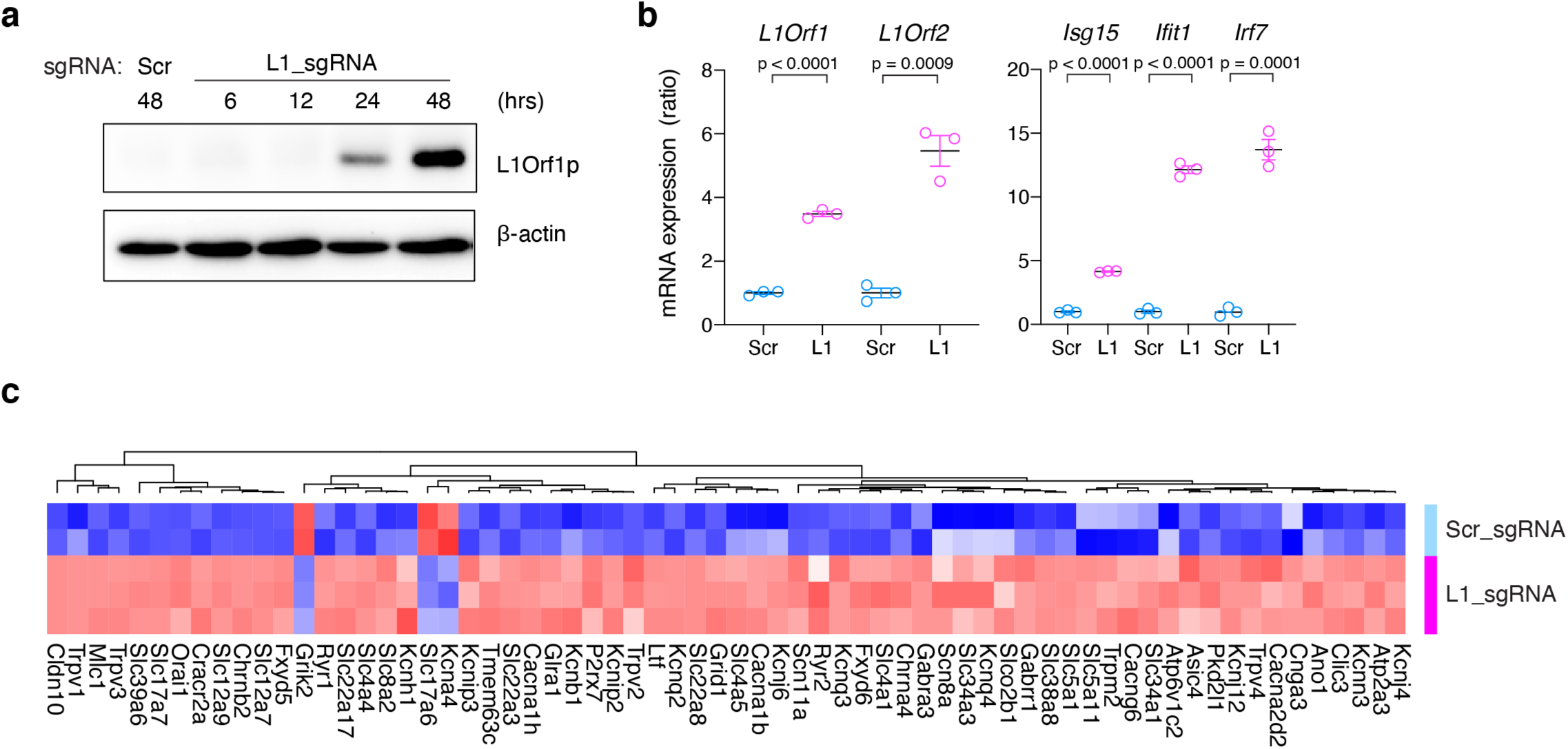
LINE-1 activation in NIH-3T3 cells in vitro. **a**, Whole cell lysates of dCasVP-3T3 cells transfected with Scr_sgRNA or L1_sgRNA expression plasmids were collected at the indicated time points after the transfection and assayed for LINE-1Orf1p detection. **b**, mRNA of dCasVP-3T3 cells transfected with Scr_sgRNA or L1_sgRNA expression plasmid (48-hour post-transfection) were analyzed with qRT-PCR (n =3). Expression levels were normalized to *Gapdh* expression and shown as the ratios to scramble condition. Data are mean ± SEM. p-values with unpaired two-sided t-test are shown. **c,** Heatmap of DEGs between L1_sgRNA and scr_sgRNA conditions in the “ion transport” term in Fig. 2b (second from the top).

**Extended Data Fig. 3.**
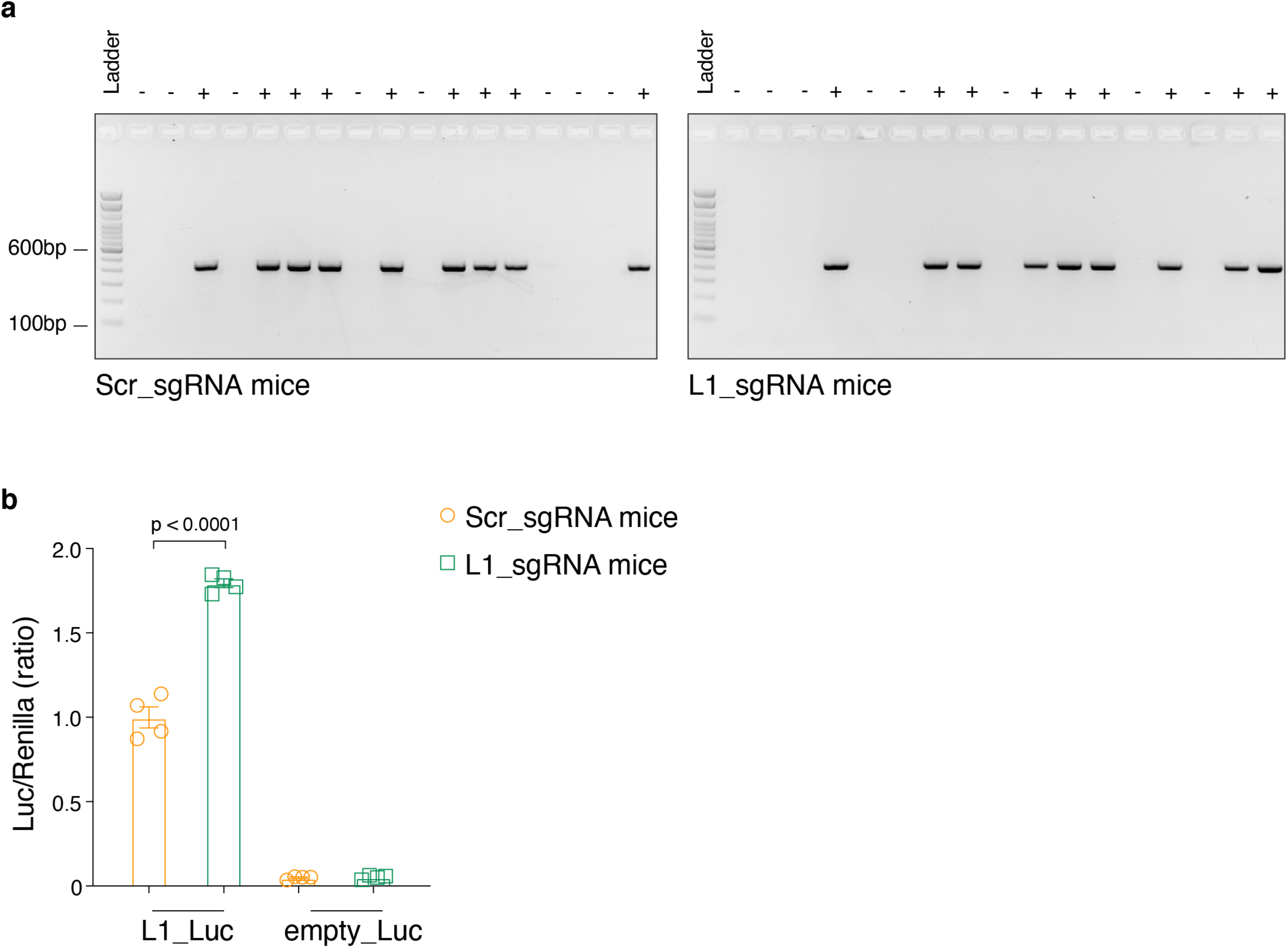
Generation of Scr_sgRNA mice and L1_sgRNA mice. **a**, Genotyping of Scr_sgRNA and L1_sgRNA mice. “+” indicates positive mice. Expected band size for both strain is 424 bp. **b**, MEFs were established from Scr_sgRNA mice and L1_sgRNA mice. CRISPRa (dCas9-VP64 and MPH) machineries were transduced in these MEFs using lentivectors, L1_Luc reporter vector or empty Luciferase vector and renilla-expression plasmid were transfected, and luciferase activities were measured. Luciferase activities were normalized to renilla activities and relative ratios to the L1_Luc+Scr_sgRNA MEF condition are shown. Data are mean ± SEM. p-value shown is with unpaired two-sided t-test.

**Extended Data Fig. 4.**
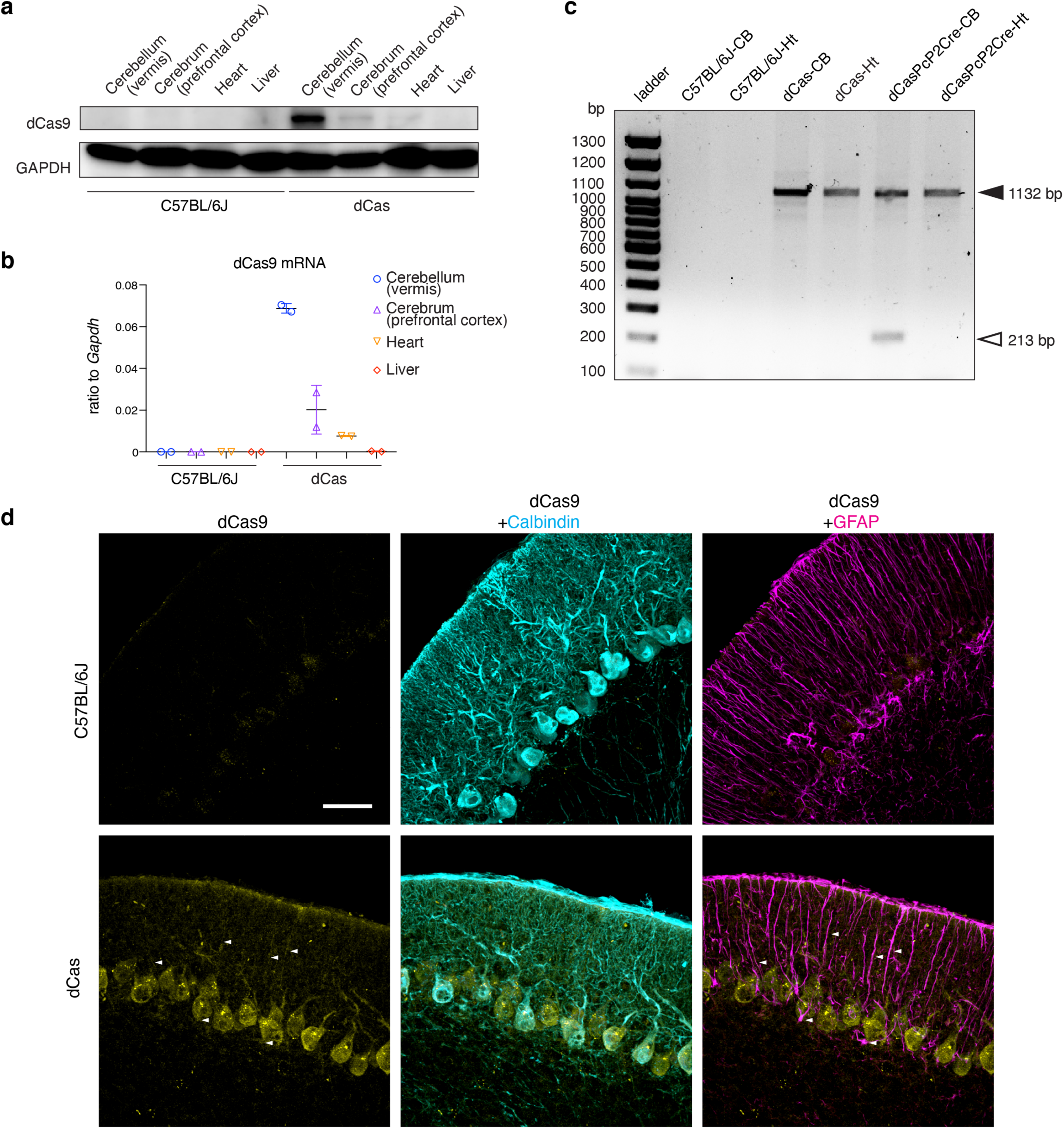
Basal expression of dCas9 in the cerebellum of dCas mice. **a**, Tissues from 8-wk-old C57/BL6J mice and dCas mice were harvested, homogenized, lysed, and the lysates were subject to Western blotting. Representative results are shown here. (n=2 mice for each group). Similar results were obtained in two independent experiments. **b**, Tissues from 8-wk-old C57/BL6J mice and dCas mice were harvested, homogenized, and mRNA was extracted. (d)Cas9 mRNA expression was assessed with qRT-PCR using specific primers for Cas9 in indicated organs. (n=2 for each group). **c**, Genomic DNAs extracted from the cerebellum (CB) and heart (Ht) from C57BL/6J, dCas, and dCas-PcP2Cre (Purkinje cell-specific Cre) double transgenic mice were subject to PCR with primers flanking LSL sequence in the dCas mice transgene. Expected band sizes for the amplification product of non-recombined and recombined sequence are 1132 bp and 213 bp, respectively (black and white arrow head, respectively). Only cerebellum of dCasPcP2Cre mice has the 213 bp product. **d**, Frozen sections of cerebella from 8-week-old dCas mice and WT (C57BL/6) mice were stained for (d)Cas9 using specific antibodies, and co-stained for Calbindin and GFAP. Representative images from n=3 for each strain are shown. Scale bar = 30 μm.

**Extended Data Fig. 5.**
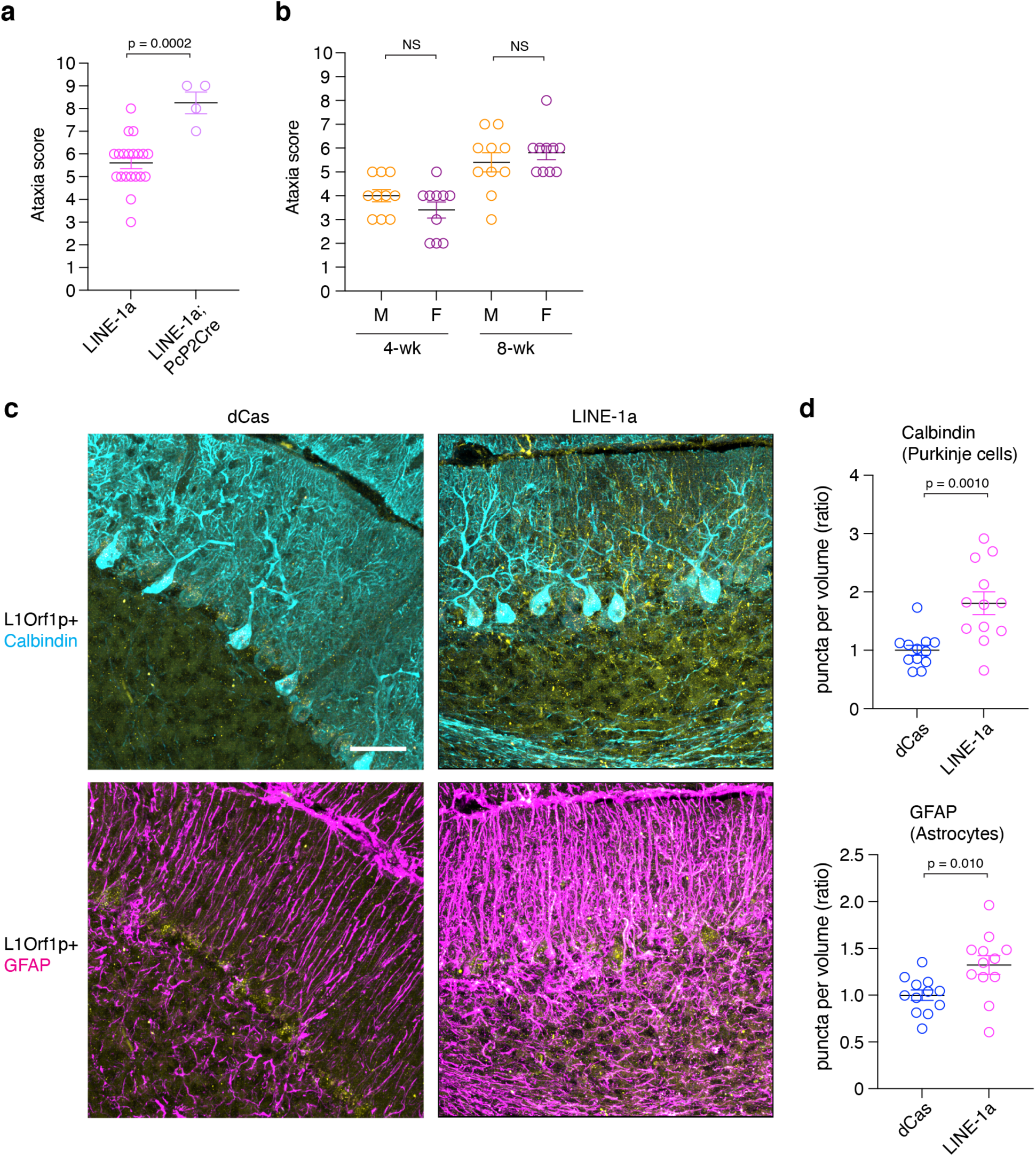
Characterization and staining of 8-wk-old LINE-1a mice. **a,** Ataxia scores were compared between 8-wk-old LINE-1a mice (n = 20, same as in Fig. 3c) and LINE-1a; PcP2Cre triple transgenic mice at the same age (transgenic for dCas9-CRISPRa, L1_sgRNA, and PcP2Cre, n = 4). **b**, Ataxia scores for LINE-1a males (M, n=10) and females (F, n=10) were compared at 4- and 8-wk of age. **c**, Representative staining images in dCas and LINE-1a mice cerebellum for LINE-1Orf1p, Calbindin, and GFAP are shown. Scale bar = 30 μm. **d**, L1Orf1p puncta densities at 8-wk-old were calculated and compared in Purkinje cells (top) and astrocytes (bottom). n = 4 per group, assessed in 3 different lobules (n = 12). **e**, GFAP staining intensities in astrocytes were compared. **a-e**, Data are mean ± SEM. p-values shown are with unpaired two-sided t-test.

**Extended Data Fig. 6.**
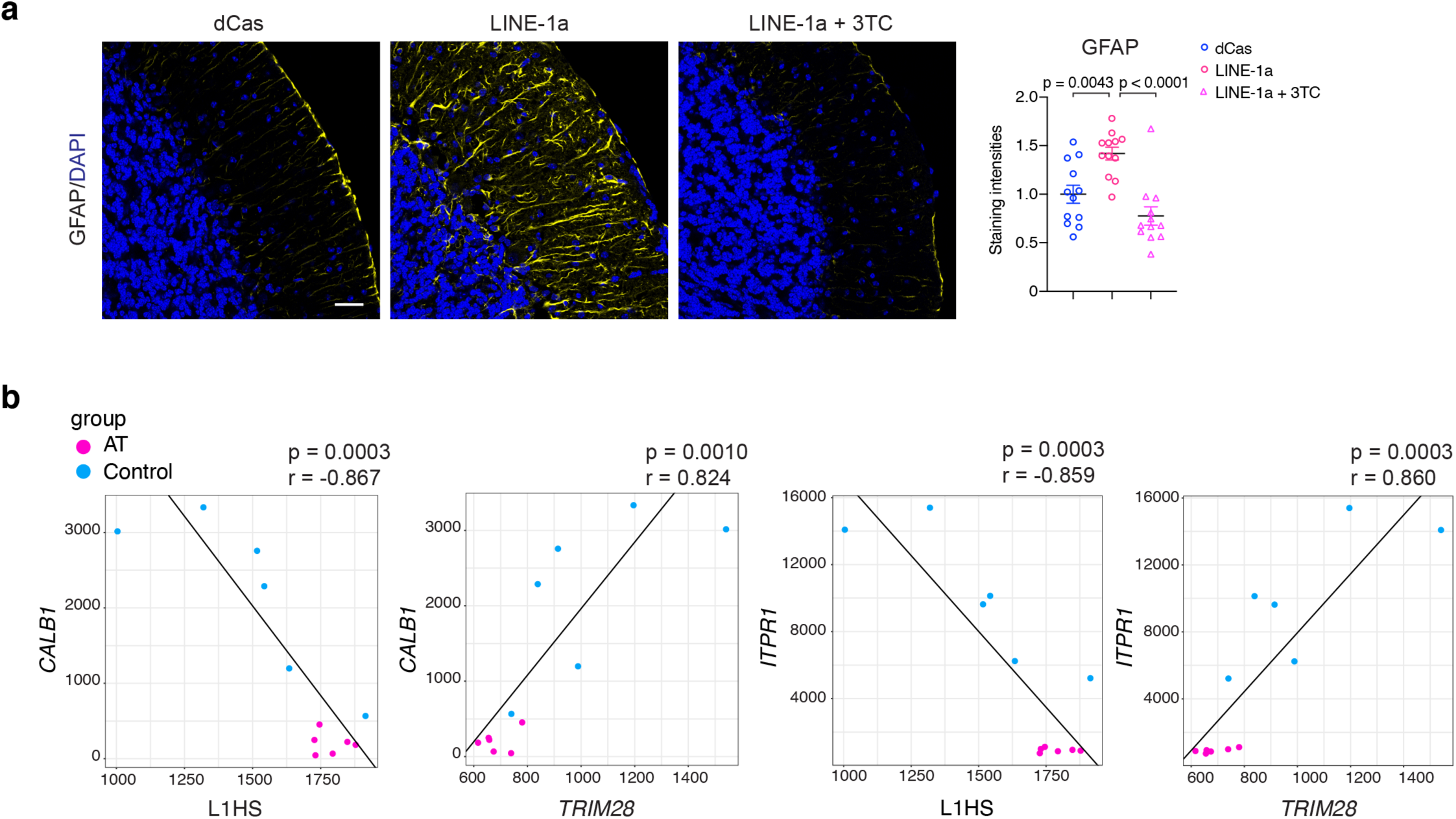
Attenuation of astrogliosis with 3TC treatment in LINE-1a mice and correlations between L1HS, *TRIM28*, *CALB1,* and *ITPR1* in human RNA-seq data. **a**, Representative images of GFAP staining in 8-wk-old dCas, LINE-1a, and 3TC-treated LINE-1a mice. Scale bar = 30 μm. Data are mean ± SEM. p-values with one-way Anova with Tukey’s post-hoc test. **b**, Correlations between expression (normalized counts) of L1HS, *TRIM28*, *CALB1*, and *ITPR1* in RNA-seq data of AT patients and control individuals (same datasets as in Fig. 1). Pearson correlation coefficients and p-values are shown.

**Extended Data Table 1.** Sample information of 33 individuals with cerebellar ataxia or unaffected control

| Clinical Brain Diagnosis | Age (yrs) | Ethnicity | Manner of Death | PMI (hrs) | Race | Sex | RNAseq |
| --- | --- | --- | --- | --- | --- | --- | --- |
| Unaffected Control | 30 | Unknown | Accidental | 20 | White | M | Yes |
| Unaffected Control | 44 | Unknown | Natural | 16 | White | F |  |
| Unaffected Control | 35 | Unknown | Natural | 17 | White | M | Yes |
| Unaffected Control | 56 | Unknown | Natural | 5 | White | M |  |
| Unaffected Control | 27 | Unknown | Natural | 13 | White | M |  |
| Unaffected Control | 25 | Unknown | Natural | 4 | White | F | Yes |
| Unaffected Control | 18 | Unknown | Natural | 22 | White | F | Yes |
| Unaffected Control | 29 | Unknown | Natural | 8 | White | M | Yes |
| Unaffected Control | 29 | Unknown | Accidental | 23 | White | F |  |
| Unaffected Control | 35 | Unknown | Natural | 27 | Black or African-American | M |  |
| Unaffected Control | 42 | Unknown | Natural | 12 | White | F |  |
| Unaffected Control | 45 | Unknown | Natural | 19 | White | M |  |
| Unaffected Control | 27 | Unknown | Natural | 12 | White | M | Yes |
| Unaffected Control | 32 | Unknown | Natural | 8 | White | F |  |
| SCA3 | 56 | Not Hispanic or Latino | Natural | 3 | White | F |  |
| SCA3 | 59 | Not Hispanic or Latino | Undetermined | 18.5 | White | M |  |
| SCA3 | 41 | Not Hispanic or Latino | Natural | 19 | White | M |  |
| SCA3 | 36 | Not Hispanic or Latino | Undetermined | 36 | White | M |  |
| SCA3 | 63 | Not Hispanic or Latino | Natural | 18.9 | White | M |  |
| SCA3 | 37 | Not Hispanic or Latino | Natural | 11.6 | White | F |  |
| Friedreich's ataxia | 27 | Unknown | Natural | 7 | White | F |  |
| Friedreich's ataxia | 36 | Not Hispanic or Latino | Natural | 42 | White | F |  |
| Friedreich's ataxia | 24 | Hispanic or Latino | Natural | 0 | Unknown | M |  |
| Ataxia Telangiectasia | 28 | Unknown | Natural | 4 | White | F |  |
| Ataxia Telangiectasia | 24 | Unknown | Natural | 2 | White | F | Yes |
| Ataxia Telangiectasia | 29 | Unknown | Natural | 12 | White | M |  |
| Ataxia Telangiectasia | 25 | Unknown | Natural | 15 | White | M | Yes |
| Ataxia Telangiectasia | 44 | Unknown | ATLL | 29 | White | M |  |
| Ataxia Telangiectasia | 27 | Not reported | Natural | 14 | White | M | Yes |
| Ataxia Telangiectasia | 36 | Unknown | Natural | 20 | White | M | Yes |
| Ataxia Telangiectasia | 27 | Not reported | Natural | 18 | White | M | Yes |
| Ataxia Telangiectasia | 31 | Not Hispanic or Latino | Natural | 4 | White | F |  |
| Ataxia Telangiectasia | 16 | Unknown | Natural | 23 | Unknown | F | Yes |

**Extended Data Table2.**
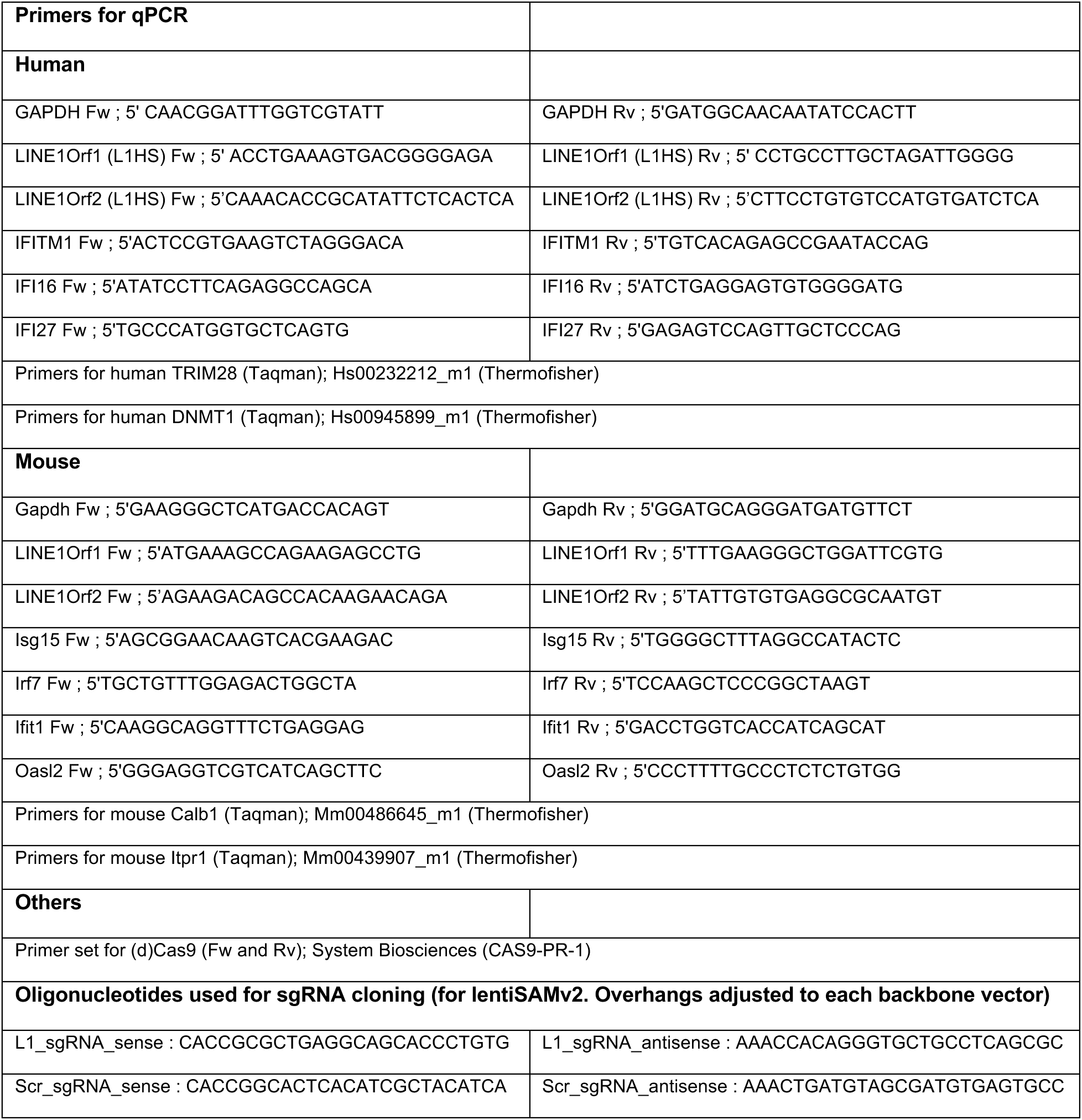
Primer and oligo sequences used in this study.

## REFERENCES

1. Saleh, A., Macia, A. & Muotri, A. R. Transposable Elements, Inflammation, and Neurological Disease. Front Neurol 10, 894, doi:10.3389/fneur.2019.00894 (2019).

2. Gorbunova, V. et al. The role of retrotransposable elements in ageing and age-associated diseases. Nature 596, 43–53, doi:10.1038/s41586-021-03542-y (2021).

3. Jonsson, M. E., Garza, R., Johansson, P. A. & Jakobsson, J. Transposable Elements: A Common Feature of Neurodevelopmental and Neurodegenerative Disorders. Trends Genet 36, 610–623, doi:10.1016/j.tig.2020.05.004 (2020).

4. Kazazian, H. H., Jr. & Moran, J. V. Mobile DNA in Health and Disease. N Engl J Med 377, 361–370, doi:10.1056/NEJMra1510092 (2017).

5. Playfoot, C. J. et al. Transposable elements and their KZFP controllers are drivers of transcriptional innovation in the developing human brain. Genome Res 31, 1531–1545, doi:10.1101/gr.275133.120 (2021).

6. Robbez-Masson, L. et al. The HUSH complex cooperates with TRIM28 to repress young retrotransposons and new genes. Genome Res 28, 836–845, doi:10.1101/gr.228171.117 (2018).

7. Castro-Diaz, N. et al. Evolutionally dynamic L1 regulation in embryonic stem cells. Genes Dev 28, 1397–1409, doi:10.1101/gad.241661.114 (2014).

8. Thomas, C. A. et al. Modeling of TREX1-Dependent Autoimmune Disease using Human Stem Cells Highlights L1 Accumulation as a Source of Neuroinflammation. Cell Stem Cell 21, 319–331 e318, doi:10.1016/j.stem.2017.07.009 (2017).

9. Hisatsune, C. & Mikoshiba, K. IP3 receptor mutations and brain diseases in human and rodents. J Neurochem 141, 790–807, doi:10.1111/jnc.13991 (2017).

10. Bezprozvanny, I. Calcium signaling and neurodegenerative diseases. Trends Mol Med 15, 89–100, doi:10.1016/j.molmed.2009.01.001 (2009).

11. Sookdeo, A., Hepp, C. M., McClure, M. A. & Boissinot, S. Revisiting the evolution of mouse LINE-1 in the genomic era. Mob DNA 4, 3, doi:10.1186/1759-8753-4-3 (2013).

12. Naas, T. P. et al. An actively retrotransposing, novel subfamily of mouse L1 elements. EMBO J 17, 590–597, doi:10.1093/emboj/17.2.590 (1998).

13. Konermann, S. et al. Genome-scale transcriptional activation by an engineered CRISPR-Cas9 complex. Nature 517, 583–588, doi:10.1038/nature14136 (2015).

14. Chen, X., Cao, R. & Zhong, W. Host Calcium Channels and Pumps in Viral Infections. Cells 9, doi:10.3390/cells9010094 (2019).

15. De Cecco, M. et al. L1 drives IFN in senescent cells and promotes age-associated inflammation. Nature 566, 73–78, doi:10.1038/s41586-018-0784-9 (2019).

16. Simon, M. et al. LINE1 Derepression in Aged Wild-Type and SIRT6-Deficient Mice Drives Inflammation. Cell Metab 29, 871–885 e875, doi:10.1016/j.cmet.2019.02.014 (2019).

17. Gasior, S. L., Wakeman, T. P., Xu, B. & Deininger, P. L. The human LINE-1 retrotransposon creates DNA double-strand breaks. J Mol Biol 357, 1383–1393, doi:10.1016/j.jmb.2006.01.089 (2006).

18. Zhou, H. et al. In vivo simultaneous transcriptional activation of multiple genes in the brain using CRISPR-dCas9-activator transgenic mice. Nat Neurosci 21, 440–446, doi:10.1038/s41593-017-0060-6 (2018).

19. Cheung, A. F., Dupage, M. J., Dong, H. K., Chen, J. & Jacks, T. Regulated expression of a tumor-associated antigen reveals multiple levels of T-cell tolerance in a mouse model of lung cancer. Cancer Res 68, 9459–9468, doi:10.1158/0008-5472.CAN-08-2634 (2008).

20. Sohal, V. S., Zhang, F., Yizhar, O. & Deisseroth, K. Parvalbumin neurons and gamma rhythms enhance cortical circuit performance. Nature 459, 698–702, doi:10.1038/nature07991 (2009).

21. Kuljis, R. O., Xu, Y., Aguila, M. C. & Baltimore, D. Degeneration of neurons, synapses, and neuropil and glial activation in a murine Atm knockout model of ataxia-telangiectasia. Proc Natl Acad Sci U S A 94, 12688–12693, doi:10.1073/pnas.94.23.12688 (1997).

22. Schwaller, B. Cytosolic Ca2+ buffers. Cold Spring Harb Perspect Biol 2, a004051, doi:10.1101/cshperspect.a004051 (2010).

23. Mikoshiba, K. IP3 receptor/Ca2+ channel: from discovery to new signaling concepts. J Neurochem 102, 1426–1446, doi:10.1111/j.1471-4159.2007.04825.x (2007).

24. Coufal, N. G. et al. Ataxia telangiectasia mutated (ATM) modulates long interspersed element-1 (L1) retrotransposition in human neural stem cells. Proc Natl Acad Sci U S A 108, 20382–20387, doi:10.1073/pnas.1100273108 (2011).

25. Jacob-Hirsch, J. et al. Whole-genome sequencing reveals principles of brain retrotransposition in neurodevelopmental disorders. Cell Res 28, 187–203, doi:10.1038/cr.2018.8 (2018).

26. Ron, D. & Walter, P. Signal integration in the endoplasmic reticulum unfolded protein response. Nat Rev Mol Cell Biol 8, 519–529, doi:10.1038/nrm2199 (2007).

27. Hetz, C., Zhang, K. & Kaufman, R. J. Mechanisms, regulation and functions of the unfolded protein response. Nat Rev Mol Cell Biol 21, 421–438, doi:10.1038/s41580-020-0250-z (2020).

28. Pereira, G. C. et al. Properties of LINE-1 proteins and repeat element expression in the context of amyotrophic lateral sclerosis. Mob DNA 9, 35, doi:10.1186/s13100-018-0138-z (2018).

29. Liu, E. Y. et al. Loss of Nuclear TDP-43 Is Associated with Decondensation of LINE Retrotransposons. Cell Rep 27, 1409–1421 e1406, doi:10.1016/j.celrep.2019.04.003 (2019).

30. Feschotte, C. Transposable elements and the evolution of regulatory networks. Nat Rev Genet 9, 397–405, doi:10.1038/nrg2337 (2008).

31. Jonsson, M. E. et al. Activation of neuronal genes via LINE-1 elements upon global DNA demethylation in human neural progenitors. Nat Commun 10, 3182, doi:10.1038/s41467-019-11150-8 (2019).

32. Sun, W., Samimi, H., Gamez, M., Zare, H. & Frost, B. Pathogenic tau-induced piRNA depletion promotes neuronal death through transposable element dysregulation in neurodegenerative tauopathies. Nat Neurosci 21, 1038–1048, doi:10.1038/s41593-018-0194-1 (2018).

33. Joung, J. et al. Genome-scale CRISPR-Cas9 knockout and transcriptional activation screening. Nat Protoc 12, 828–863, doi:10.1038/nprot.2017.016 (2017).

34. Cong, L. et al. Multiplex genome engineering using CRISPR/Cas systems. Science 339, 819–823, doi:10.1126/science.1231143 (2013).

35. Dobin, A. et al. STAR: ultrafast universal RNA-seq aligner. Bioinformatics 29, 15–21, doi:10.1093/bioinformatics/bts635 (2013).

36. Treger, R. S. et al. The Lupus Susceptibility Locus Sgp3 Encodes the Suppressor of Endogenous Retrovirus Expression SNERV. Immunity 50, 334–347 e339, doi:10.1016/j.immuni.2018.12.022 (2019).

37. Tokuyama, M. et al. ERVmap analysis reveals genome-wide transcription of human endogenous retroviruses. Proc Natl Acad Sci U S A 115, 12565–12572, doi:10.1073/pnas.1814589115 (2018).

38. Criscione, S. W., Zhang, Y., Thompson, W., Sedivy, J. M. & Neretti, N. Transcriptional landscape of repetitive elements in normal and cancer human cells. BMC Genomics 15, 583, doi:10.1186/1471-2164-15-583 (2014).

39. Quinlan, A. R. & Hall, I. M. BEDTools: a flexible suite of utilities for comparing genomic features. Bioinformatics 26, 841–842, doi:10.1093/bioinformatics/btq033 (2010).

40. Love, M. I., Huber, W. & Anders, S. Moderated estimation of fold change and dispersion for RNA-seq data with DESeq2. Genome Biol 15, 550, doi:10.1186/s13059-014-0550-8 (2014).

41. Huang da, W., Sherman, B. T. & Lempicki, R. A. Systematic and integrative analysis of large gene lists using DAVID bioinformatics resources. Nat Protoc 4, 44–57, doi:10.1038/nprot.2008.211 (2009).

42. Sulkowski, P. L. et al. 2-Hydroxyglutarate produced by neomorphic IDH mutations suppresses homologous recombination and induces PARP inhibitor sensitivity. Sci Transl Med 9, doi:10.1126/scitranslmed.aal2463 (2017).

43. Sulkowski, P. L. et al. Krebs-cycle-deficient hereditary cancer syndromes are defined by defects in homologous-recombination DNA repair. Nat Genet 50, 1086–1092, doi:10.1038/s41588-018-0170-4 (2018).

44. McQuin, C. et al. CellProfiler 3.0: Next-generation image processing for biology. PLoS Biol 16, e2005970, doi:10.1371/journal.pbio.2005970 (2018).

45. Guyenet, S. J. et al. A simple composite phenotype scoring system for evaluating mouse models of cerebellar ataxia. J Vis Exp, doi:10.3791/1787 (2010).

46. Liu, Z. W. & Gao, X. B. Adenosine inhibits activity of hypocretin/orexin neurons by the A1 receptor in the lateral hypothalamus: a possible sleep-promoting effect. J Neurophysiol 97, 837–848, doi:10.1152/jn.00873.2006 (2007).

47. Liu, Z. W., Gan, G., Suyama, S. & Gao, X. B. Intracellular energy status regulates activity in hypocretin/orexin neurones: a link between energy and behavioural states. J Physiol 589, 4157–4166, doi:10.1113/jphysiol.2011.212514 (2011).

48. Franklin, K. B. J. & Paxinos, G. Paxinos and Franklin’s The mouse brain in stereotaxic coordinates. Fourth edition. edn, (Academic Press, an imprint of Elsevier, 2013).

49. Somogyi, P. & Takagi, H. A note on the use of picric acid-paraformaldehyde-glutaraldehyde fixative for correlated light and electron microscopic immunocytochemistry. Neuroscience 7, 1779–1783, doi:10.1016/0306-4522(82)90035-5 (1982).

50. Tokuyama, M. et al. Antibodies against human endogenous retrovirus K102 envelope activate neutrophils in systemic lupus erythematosus. J Exp Med 218, doi:10.1084/jem.20191766 (2021).

